# Global analysis of aging-related protein structural changes uncovers enzyme polymerization-based control of longevity

**DOI:** 10.1101/2023.01.23.524173

**Authors:** Jurgita Paukštytė, Rosa María López Cabezas, Yuehan Feng, Kai Tong, Daniela Schnyder, Ellinoora Elomaa, Pavlina Gregorova, Matteo Doudin, Meeri Särkkä, Jesse Sarameri, Alice Lippi, Helena Vihinen, Juhana Juutila, Anni Nieminen, Petri Törönen, Liisa Holm, Eija Jokitalo, Anita Krisko, Juha Huiskonen, L. Peter Sarin, Ville Hietakangas, Paola Picotti, Yves Barral, Juha Saarikangas

## Abstract

Aging is associated with progressive phenotypic changes over time. Virtually all cellular phenotypes are produced by proteins and structural alterations in proteins can lead to age-related diseases. Nonetheless, comprehensive knowledge of proteins undergoing structural-functional changes during cellular aging and their contribution to age-related phenotypes is lacking. Here, we conducted proteome-wide analysis of early age-related protein structural changes in budding yeast using limited proteolysis-mass spectrometry. The results, compiled in online ProtAge-catalog, unravelled age-related functional changes in regulators of translation, protein folding and amino acid metabolism. Mechanistically, we found that folded glutamate synthase Glt1 polymerizes into supramolecular self-assemblies during aging causing breakdown of cellular amino acid homeostasis. Inhibiting Glt1 polymerization by mutating the polymerization interface restored amino acid levels in aged cells, attenuated mitochondrial dysfunction and led to life span extension. Altogether, this comprehensive map of protein structural changes enables identifying novel mechanisms of age-related phenotypes and offers opportunities for their reversal.

## Introduction

The three-dimensional structures of proteins define how genetic information is translated into cellular biochemistries. As such, structural information of proteins is required to fully comprehend how cellular and organismal phenotypes arise ^1–4^. Most proteins can occupy various structural states in different contexts. Variation can result, for example, from interactions with other molecules ^5,6^, post-translational modifications ^7^, fold switching ^8^, unstructured regions ^9,10^ and condensation-reactions ^11,12^. However, at present we know little of the diversity and dynamics of protein structural variation and even less about how it contributes to phenotypic diversity of cells and organisms.

Aging is a nearly universal biological process characterized by progressive phenotypic changes and increased mortality over time ^13,14^. The rate of these changes is influenced by genetic variance, life history and environment. These variables can be experimentally controlled in model organisms, allowing for a more precise investigation of the mechanisms underlying aging phenotypes. For instance, the single-celled organism *Saccharomyces cerevisiae* starts to systematically display aging phenotypes during early stages of replicative aging that are not caused by mutations ^15–17^. Such phenotypes include evolutionarily conserved aging hallmarks such as changes in cellular size ^18^, metabolism ^19,20^, organelle function ^21–23^ and the formation of protein aggregates ^24,25^. Understanding how these and other phenotypic changes in aging arise can lead to the discovery of early causal mechanisms of cascading negative effects and thereby provide opportunities for interventions.

It is still not clear what gives rise to progressive phenotypic changes in aging. The prevailing theory is that molecular damage, for example due to free radicals ^13,26^ in conjunction with changes in the fidelity of epigenetic modifications ^27^, transcription ^28,29^, translation ^30–32^ and metabolism ^19^, contributes to physiological changes in aging. In addition, aging is associated with alterations in the abundance, turnover or folding of proteins that are typically linked to defective protein homeostasis ^33–35^. Folding changes in turn can lead to protein aggregation, which is a shared feature of aging across organisms and tissue types ^36,37^, and is infamously linked to age-related neurodegenerative diseases ^38^. However, it is currently unclear whether cellular aging results in other types of protein structural changes beyond aggregation. Thus, a more comprehensive characterization of protein structural states is required to understand the full breadth of proteome activity changes during aging and their contribution to adaptive and maladaptive age-related phenotypes.

## Results

### Global analysis of aging-associated structural changes

To identify aging-associated structural changes on a proteome-wide scale, we used magnetic-activated affinity purification to obtain fractions of young (av. age 0.6 divisions ±0.91 STD) and aged (av. age 4.2 divisions ±0.65 STD) budding yeast cells (**Fig. 1A**). Yeast cells aged for an average of 4.2 divisions represents mother cells that begin to systematically display aging phenotypes that are not caused by mutations ^15–17^, enabling us to capture the underlying processes behind these age-related changes as they emerge. Such phenotypes include conserved aging hallmarks such as the appearance of protein aggregates and the declined function of organelles such as mitochondria and vacuoles ^21,22,25^. The young cells represent rejuvenated daughter cells (since aging factors are retained in the mother cell) ^15–17,39^. We extracted native proteomes from young and aged cells, subjected them to limited proteolysis (LiP) via brief proteinase K treatment, followed by denaturation, trypsin digestion and mass spectrometry analysis (LiP-MS) ^40,41^. A consequence of LiP is that protein structural features impact the cleavage patterns, enabling identification of age-related structural alterations in proteins at peptide-level resolution (**Fig. 1A**). To normalize the LiP-MS results for age-related changes in protein abundance, the same samples were also analyzed without the LiP step.

**Figure 1.**
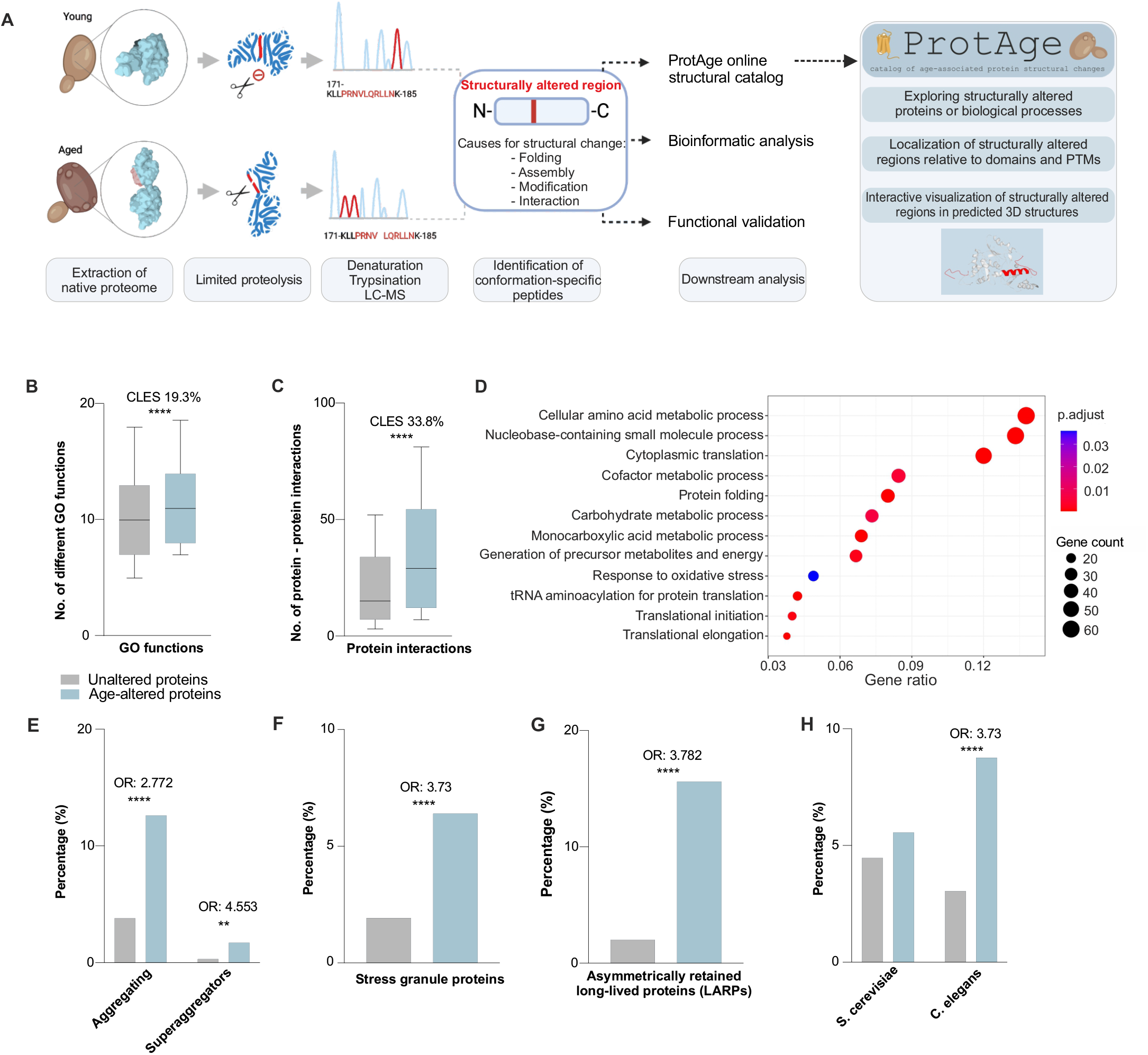
Identification of age-related structural changes in pleiotropic regulators of cell metabolism, growth, and stress responses. **A**. Overview of workflow to identify age-related protein structural changes using LiP-MS. **B**. Number of GO functions in age-altered proteins (light blue) compared to unaltered proteins (grey). **C**. Number of protein-protein interactions in age-altered proteins (*n* = 460) (light blue) compared to unaltered proteins (*n* = 2279) (grey). Black line in box plots in (B-C) is median and whiskers mark 10 and 90 percentile values. **D**. Gene ontology (GO) enrichment by over-representation analysis of age-altered proteins compared to unaltered proteins. The size of dots in the plot represents the number of proteins for every biological process. The color change from blue to red signify the size of adjusted *P* value (*P* adjusted < 0.05). **E**. Fraction of aggregating proteins and superaggregators among age-altered proteins (*n* = 468) (light blue) and unaltered proteins (*n* = 2332) (grey). **F**. Fraction of stress granules forming proteins among age-altered proteins (*n* = 468) (light blue) compared to unaltered proteins (*n* = 2332) (grey). **G**. Fraction of asymmetrically retained long-lived proteins (LARPs) among age-altered proteins (*n* = 468) (light blue) compared to unaltered proteins (*n* = 2332) (grey). **H**. Fraction of genes whose decreased expression extends lifespan in age-altered (*n* = 468) (light blue) and unaltered proteins (*n* = 2332) (grey) or their homologs in *S. cerevisiae* and *C. elegans*. Statistical significance was assessed with Fisher’s exact test and effect size in (B-C) is displayed as common language effect size (CLES) and in (E-H) as odds ratio (OR). *\*\*P* < 0.01; *\*\*\*\*P* < 0.0001.

We considered that proteins with significant differences in proteolytic cleavage patterns normalized to their abundance between young and aged cells have undergone an aging-related structural change (collectively referred to hereafter as *age-altered proteins*) (**Fig. 1A**). LiP-MS enables reproducible identification of structural changes caused for example by alterations in protein folding, assembly state, interaction with other molecules or post-translational modifications, thereby offering a readout for age-associated changes in protein function ^3,40,42^.

The global comparison of protein structural states between young and aged cells uncovered age-dependent structural differences in 468 proteins that comprised 1272 conformation-specific peptides (fold change q-value < 0.05 from sample triplicates, total proteins detected: 2833) (**Fig. 1A**). These conformotypic peptides identify the protein regions that changed structure during early aging. We mapped all structurally altered regions to their Alpha Fold-predicted three dimensional (3D) protein structures ^43^ and collected these results together with other relevant features into the online ProtAge-catalog (https://protage-server-21.it.helsinki.fi/) (**Fig. S1A)**.

To uncover shared features of age-altered proteins, we compared the structurally altered proteome to the detected proteome that did not show significant structural changes (*unaltered proteins*). We found that age-altered proteins had on average more assigned gene ontology (GO) terms and protein-protein interactions, suggesting that these proteins are pleiotropic and participate in diverse cellular functions (**Fig. 1B-C, Fig. S1B**). GO enrichment analyses showed that regulators of metabolism, translation, protein folding and oxidative stress responses were overrepresented in the age-altered proteome (p.adj < 0.05 **Fig. 1D**). The age-altered proteins contained a larger fraction of essential genes and had a lower mutation rate than the unaltered proteins (**Fig. S1C-D**). Interestingly, even though the distribution of protein solubility was similar between age-altered proteins and the unaltered proteins (**Fig. S1E**), there was a strong enrichment for proteins that can transition into biomolecular condensates during stress ^44^, including stress granule components (**Fig. 1E-F**). Moreover, age-altered proteins were strongly enriched in a class of long-lived proteins that are retained in the aging mother cells, suggesting that their retention is associated with an underlying structural change ^45^ (**Fig. 1G**). In addition to being long-lived, the age-altered proteins were more abundant than the unaltered proteins, which could be explained by the overall higher translation rate and lower number of degradation signals in these proteins (**Fig. S1F-H**). Finally, we found that decreased expression of genes encoding for the age-altered proteins in budding yeast or their homologs in *Caenorhabditis elegans* were more readily associated with life span extension as compared to the genes encoding for the unaltered proteins (**Fig. 1H**). Taken together, aging-associated structural changes are prevalent in pleiotropic regulators of metabolism, growth and stress responses that can transition between soluble and assembled states during stress.

### Structural changes identify activity changes in regulators of protein homeostasis

Can the structural changes provide new insights in protein function during aging? We observed significant structural changes in all major chaperone classes, including members of Hsp40, Hsp70, Hsp90, Hsp100, Hsp110 and TRiC/CCT proteins, which are interconnected and implicated in the maintenance of proteostasis during aging **(Fig. S2A)** ^33,34,46^. The major Hsp70 isoform Ssa1 availability is compromised during aging ^47^, but how this decline in function relates to structure is unclear. To address this question, we mapped the structurally altered regions in Ssa1. Intriguingly, they clustered around two major sites undergoing activity-dependent structural changes during the Hsp70 allosteric cycle: the nucleotide binding domain (NBD) and the substrate binding domain (SBD) ^48^ (**Fig. 2A, Fig. 2B**). Moreover, all three full tryptic peptides localizing to the SBD showed significantly increased abundance in the aged cells as compared to young cells, suggesting that this region in SBD (residues 413-447) was less accessible for proteolytic cleavage in aged cells (**Fig. 2A**). To investigate whether the reduced accessibility of SBD reflects increased occupancy by clients, we performed a contact-site analysis using the substrate bound structure of bacterial Hsp70 DnaK. The proteolytically protected regions overlapped with the evolutionarily conserved residues that mediate critical hydrogen bonding and hydrophobic interactions with the clients ^49,50^ (**Fig. 2B**). These data are consistent with the increased client-binding of Ssa1 in aged cells, as previously measured by a decreased diffusion coefficient ^47^ and increased association with protein aggregates ^25^ (**Fig. 2C**). The results therefore provide validation of the ability of the LiP-MS approach to identify relevant aging-associated proteomic changes. Interestingly, the ribosome-associated Hsp70 proteins Ssb1 and Ssb2 and the endoplasmic reticulum Kar2/BiP showed a similar pattern of structural alterations in the SBD regions that overlapped with the client-binding site ^49,50^ (**Fig. S2C-H**), suggesting that aging might also lead to increased client occupancy of Hsp70s that govern nascent protein folding in cytoplasm and ER, respectively. Overall, we propose that the observed structural changes in Hsp70 proteins reflect an overall age-related increase in client occupancy, and this change may impact the proteostasis of cytosolic and ER residing proteins.

**Figure 2:**
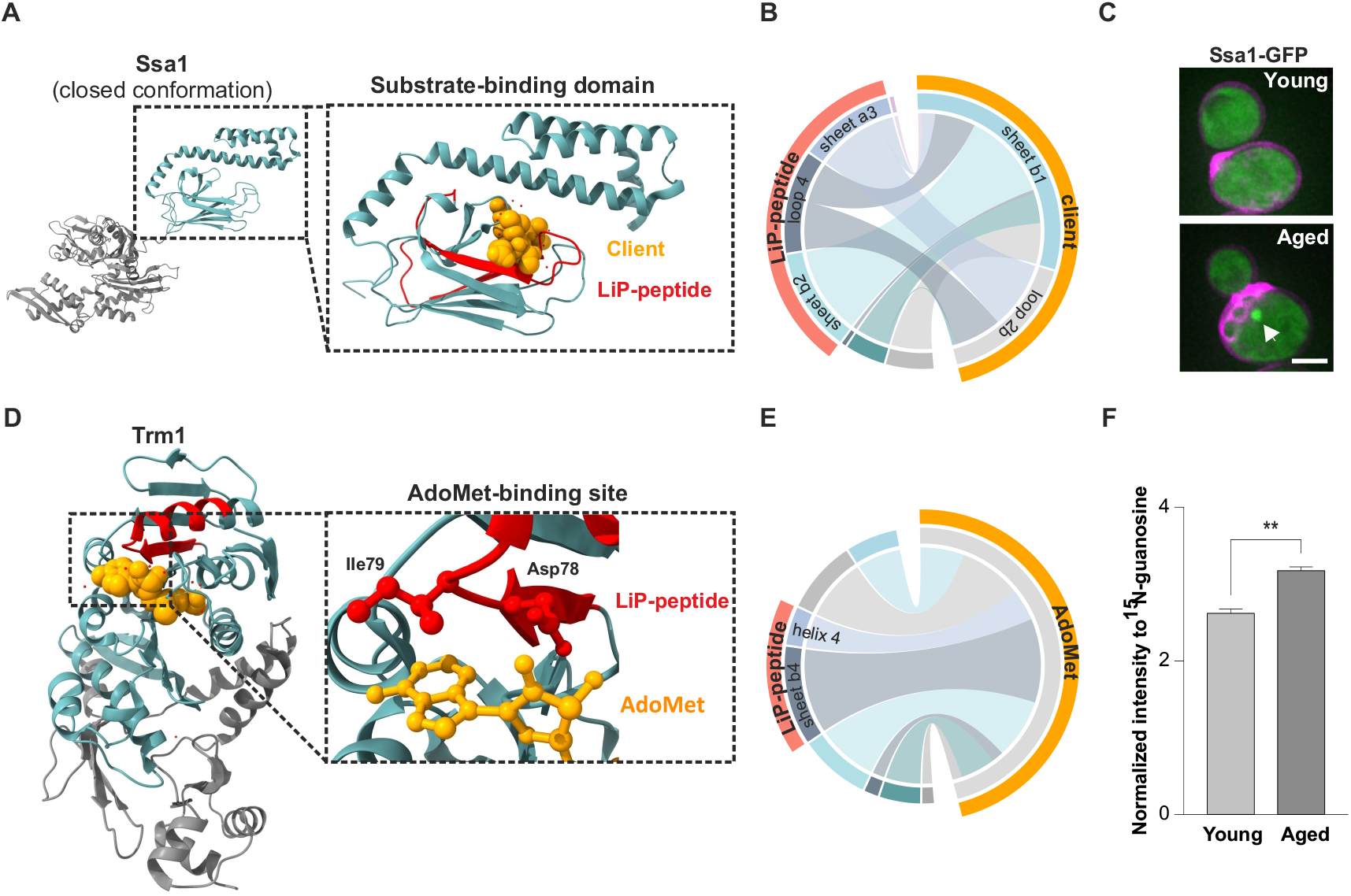
Structural alterations indicate age-related protein activity changes in regulators of proteostasis. **A**. Cartoon representation of ADP-bound ‘open’ Hsp70 with the nucleotide binding domain in gray and the substrate binding domain (SBD) in light green (PDB:2KHO; ^74^). Close-up of the SBD with Ssa1 LiP-peptides mapped in red and the peptide substrate in yellow (PDB:1DKX; ^49^). **B**. The chord plot depicts contacts between LiP peptide (red) and client peptide (orange) based on (PDB:1DKX). The inner arcs display the contacting secondary structures. **C**. Representative images of young and aged cells expressing Ssa1 tagged with GFP show association of Ssa1 with protein aggregates in aged cells (arrowhead). Cell age can be determined from bud scars on cell wall stained with calcofluor (magenta). Scale bar 2 μm. **D**. Cartoon structure of *Pyrococcus horikoshii* Trm1. LiP-peptide depicted in red and AdoMet in yellow. Close up showing two AdoMet interacting residues Asp78 and Ile79. (PDB: 2EJT; ^52^). **E**. Chord plot displays contacts between LiP peptide-region (red) and AdoMet (yellow). The inner arcs depict contacting secondary structures. **F**. Levels of N2,N2-dimethylguanosine (m^2^,^2^G) modified tRNA increase in aged (mean age 4.4 divisions) cells compared to young (mean age 0.1 divisions) cells. TRNA levels were identified by mass spectrometry and normalized to an internal ^15^N-guanosine standard. Significance was determined with two tailed unpaired t-test with Welch’s correction. *n* = 2. \*\**P* < 0.01.

One cause of compromised proteostasis during aging is altered protein translation accuracy ^31,32^. Indeed, age-altered proteins were enriched in proteins regulating translation and tRNA aminoacylation and included some tRNA modifying enzymes (**Fig. 1D**). tRNA modifications regulate the kinetics and accuracy of translating ribosomes and thereby contribute to protein folding ^51^. However, it is unknown whether specific tRNA modifications change during aging. We noticed that Trm1, which catalyzes N2,N2-dimethylguanosine (m^2^,^2^G) modification at a position G_26_ in tRNAs, undergoes an age-related structural change that localized to the conserved Motif II (**Fig. 2D)**. This motif is part of the methyl donor molecule binding pocket that is important for Trm1 catalytic activity. Contact site analysis revealed that the LiP-peptide directly overlaps with S-adenosyl-l-methionine (AdoMet) binding site and included the conserved Asp78 that mediates hydrophilic interaction with the ribose group of AdoMet ^52^ (**Fig. 2D-E, Fig. S3A**). The modification site specificity of Trm1 on tRNAs (**Fig. S3B**) enabled us to test whether the observed structural change corresponds to an activity change during aging. For this, we extracted tRNAs from young and aged cells and quantified the m^2^,^2^G modification by mass spectrometry. Interestingly, we found a significantly increased abundance of m^2^,^2^G in aged cells (**Fig. 2F**), consistent with increased occupancy of Trm1 active site in aged cells.

Overall, we found structural changes in key proteins regulating proteostasis at the level of translation and folding, consistent with age-related increased substrate association. More generally, these results underline that LiP-MS can facilitate identification of specific age-related functional changes in proteins and guide their mechanistic analysis.

### Glutamate synthase Glt1 transitions to mesoscale assemblies during aging

Amino acid metabolizing enzymes were the most overrepresented class of age-altered proteins (**Fig. 1D and SFig. 4A**). To evaluate if the structural changes impact function, we chose to examine whether age-altered metabolic enzymes show changes in subcellular localization or organization, we used fluorescence microscopy to visualize GFP-tagged versions of endogenous cytoplasmic amino acid synthesizing enzymes in young and aged cells. Most enzymes showed diffuse cytoplasmic localization with no detectable age-related changes (**Fig. S4B**). A subset of enzymes showed a punctate pattern, which in some cases appeared to be associated with localization to organelles (**Fig. S4C**). Intriguingly, unlike all other proteins in this class, the glutamate synthase Glt1 displayed a clear and systematic age-related localization change: in young cells Glt1 was diffusely distributed throughout the cytoplasm, but in aged cells that had undergone 4-6 divisions Glt1 had transitioned into large rod-shaped structures, which we show below are Glt1 polymers (**Fig. 3A-B, Fig. S4C, Movie S1**). The Glt1 polymers formed in a similar manner when tagged with either GFP or monomeric mKate2 fluorophores (**Fig. S4D**).

**Figure 3:**
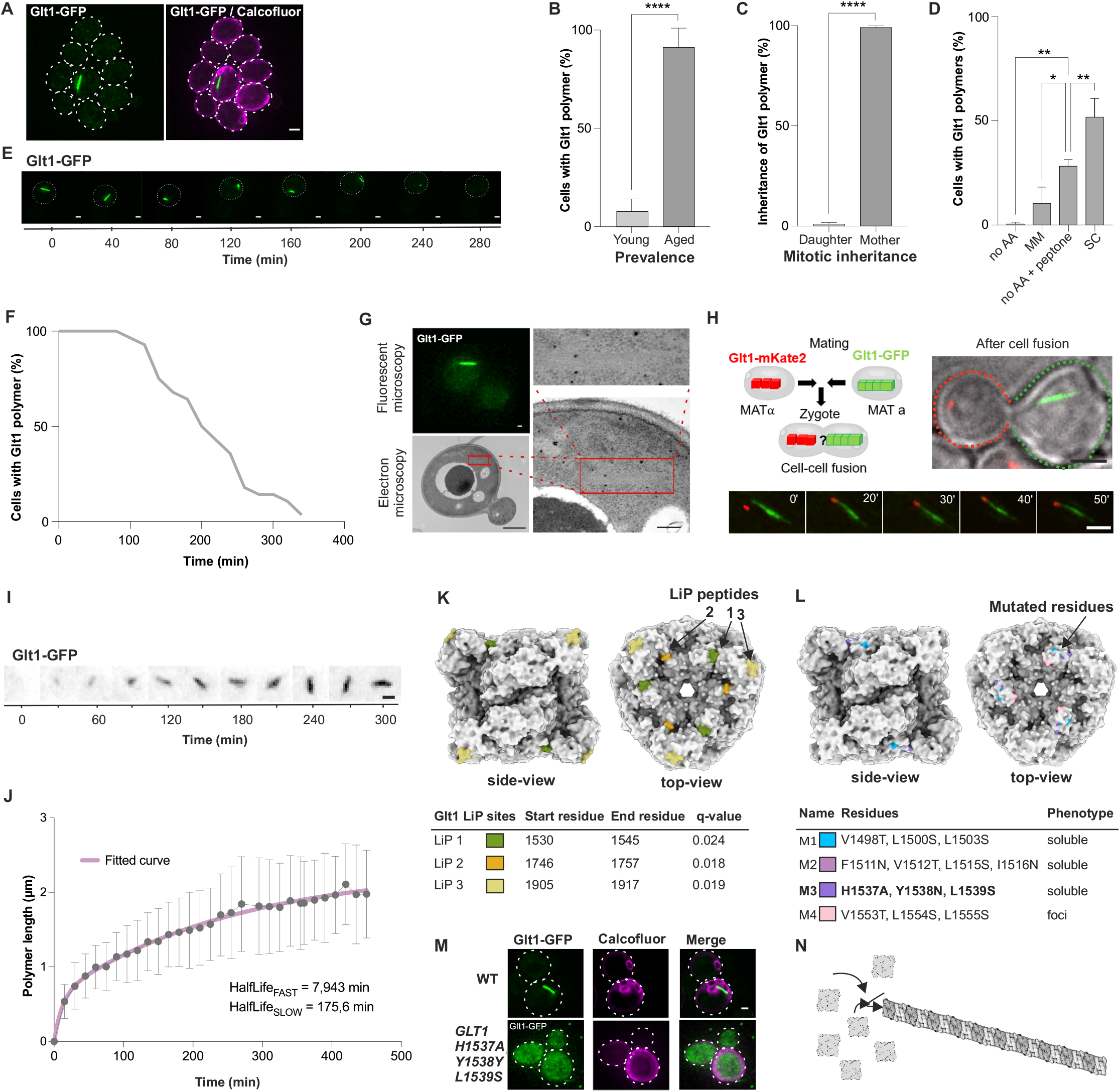
Glutamate synthase Glt1 forms filamentous polymers during aging. **A**. Endogenous Glt1 tagged with GFP is soluble in young cells but forms a fiber-like polymers in aged cells. Scale bar 1 μm. **B**. Quantification of cells bearing Glt1 polymers between young (0-1 divisions, *n* = 681) and aged (4-6 divisions, *n* = 82) cells. **C**. Quantification of inheritance of Glt1 polymers by mother or daughter cells during cell division by live-cell imaging. *n* = 364. **D**. Quantification of cells with Glt1 polymers in varying amino acid conditions: no amino acids (no AA), with peptone (no AA + peptone) and synthetic complete media (SC). *n* = 3. **E**. representative time-lapse images from microfluidics-coupled microscopy analysis of Glt1-GFP (green) polymer over time after switching to amino acid deprived media. Scale bar 1 μm. **F**. Quantification of Glt1 polymer dynamics over time after switching to media without amino acids. *n* = 40. **G**. Correlative light-electron microscopy image of Glt1-GFP (green) polymer bearing cell detected with fluorescence microscopy and subsequently visualized by transmission electron microscopy (TEM). Area of Glt1 polymer denoted with the box. Scale bar is 1 μm and in the zoom-in EM image (right) 200 nm. **H**. Mating cells carrying Glt1 polymers tagged with mKate2 (red) or GFP (green). Time lapse images of a stable co-assembly of two polymers. Scale bar 2 μm. **I**. Time-lapse microscopic evaluation of Glt1-GFP polymer assembly. Scale bar 1 μm. **J**. Dynamics of Glt1 polymer assembly was assessed by measuring the length of assemblies in individual cells over 345 min after nucleation and plotted with two-phase association model, showing duplication time for fast 7,943 min (HalfLife_FAST_) and slow 175.6 min (HalfLife_slow_) states. Red line displaying model fit (R^2^ = 0.53). *n* = 77. **K**. Table summarizing identified LiP peptides in Glt1 and visualized in a structural model of yeast Glt1 hexamer (PDB: 2VDC; ^57^). **L**. Color-coded representation of mutated residues in a Glt1 structural model. Table summarizes the effect of the different mutants on Glt1 polymerization. **M**. Representative images wild type and Glt1 H1537A, Y1538N, L1539S mutant cells tagged with GFP. Scale bar 1 μm. **N**. Cartoon model of putative polymerization mechanism of Glt1 with dihedral symmetry (D3). Error bars in the graphs represent ± SD. Statistical significance was assessed with two tailed unpaired t test (B-C) or one-way ANOVA with Dunnett’s correction. *\*P* < 0.05; *\*\*P* < 0.01, *\*\*\*\*P* < 0.0001.

**Figure 4:**
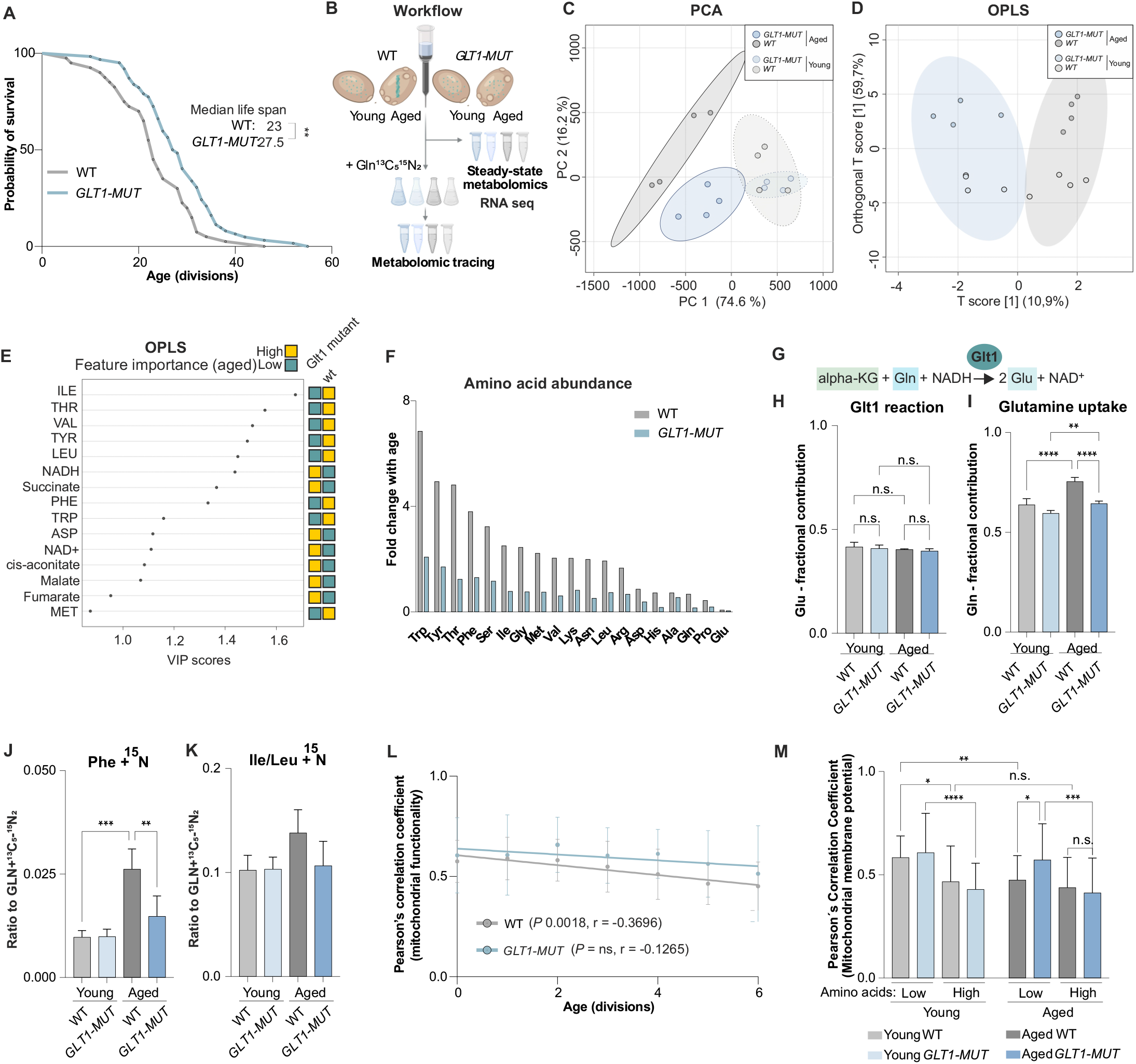
Glt1 polymerization is reversible and promotes accumulation of intracellular amino acids. **A**. Replicative life span comparison between WT and *GLT1-MUT* cells. The median life span is displayed on the graph. *n*_*wt*_ = 40, *n*_*GLT1-MUT*_ = 62. Statistical significance was assessed with Mantel-Cox test. **B**. Workflow for comparing transcriptomics and metabolomics. Purified fractions of young and aged cells were subjected to RNAseq and steady-state metabolomics analysis or grown with labelled ^13^C_5_^15^N_2_-glutamine for tracing analysis. **C**. Principal component analysis (PCA) of steady-state metabolomics. Young cell clusters encircled with dotted lines, aged cell clusters with solid lines. *n* = 4. **D**. Differences in steady-state metabolites between WT and *GLT1-MUT* are summarized in orthogonal partial least square (OPLS) analysis. *n* = 4. **E**. Feature importance graph of OPLS indicating metabolites and their weight with VIP score between aged WT and *GLT1-MUT* (yellow higher and turquoise lower metabolites levels between two groups). *n* = 4. **F**. Fold change in steady-state amino acid levels with age in WT and *GLT1-MUT* cells *n* = 4. **G**. Reaction catalyzed by Glt1. **H**. The fractional contribution of labelled glutamate from Glt1 reaction. *n* = 4. **I**. Fractional contribution of labelled ^13^C_5_-^15^N_2_-glutamine uptake. *n* = 4. **J**. Utilization of labelled ^13^C_5_-^15^N_2_-glutamine for phenylalanine (Phe) synthesis via transamination is shown by a ratio of labelled ^13^C_5_-^15^N_2_-glutamine to ^15^N-labelled Phe. *n* = 4. **K**. Utilization of labelled ^13^C_5_-^15^N_2_-glutamine for isoleucine/leucine (Ile/Leu) synthesis via transamination shown by a ratio of labelled ^13^C_5_-^15^N_2_-glutamine to ^15^N-labelled Ile/Leu. *n* = 4. **L**. Age-dependent analysis of mitochondrial membrane potential between WT and *GLT1-MUT* cells. Correlation coefficient (R) and its significance was computed with standard Pearson’s correlation test where the dots represent single cells and line indicates the linear regression curve. *n*_*WT*_ = 216, *n*_*GLT1-MUT*_ = 154. **M**. Quantification of MitoLoc signal in normal and high amino acid conditions in WT and *GLT1-MUT* cells. *n*_*WT*_ = 186, *n*_*GLT1-MUT*_ = 149, at least 25 cells per subgroup was analyzed from 3 biological replicates. Error bars represent mean ± SD. Statistical significance in (H-K) was assessed with ordinary one-way ANOVA with Dunnett’s correction and in M with three-way ANOVA with Šidák’s correction. *\*P* < 0.05; *\*\*P* < 0.01; *\*\*P* < 0.001 *\*\*\*\*P* < 0.0001; n.s. not significant.

Metabolic enzymes, including Glt1, were previously observed to form polymers in energy-starved, non-cycling cells ^53–56^. The Glt1 structures resembled these polymers. However, the aged cells carrying Glt1 polymers were cycling normally (**Movie S1**). Live-cell imaging further revealed that Glt1 polymers were retained in the aging mother cells during cell division, and the daughter cells were born with diffuse Glt1 (**Fig. 3C, Movie S1**). The timing of appearance and mitotic segregation pattern of Glt1 assemblies resembled that of aging-associated aggregates ^25,47^. However, Glt1-GFP polymers did not colocalize with Hsp104-labelled aggregates (**Fig. S4E**), indicating that Glt1 polymers do not have features of canonical aggregates.

As Glt1 polymers did not appear to be aggregates, we investigated how their formation is regulated. We tested if Glt1 polymerization responds to availability of key nutrients by modulating the levels of glucose, nitrogen and amino acids. Interestingly, we found that Glt1 polymerization was affected by amino acid availability but not by limiting glucose or nitrogen (**Fig. 3D, Fig. S5A-B**). Incubating cells in growth medium lacking amino acids inhibited Glt1 polymer formation during replicative aging, whereas the addition of minimal amino acids for growth, methionine, histidine and leucine (minimal media, MM) was sufficient to restore polymer formation (**Fig. 3D**). Polymer formation was further increased by adding peptone as the source for all amino acids, or by using synthetic complete (SC) media, that contains most amino acids (see methods for details) (**Fig. 3D**). These data suggest that Glt1 polymerization during aging is a tunable reaction that is regulated by amino acid levels.

To test if Glt1 polymers are reversible and can depolymerize, we used microfluidics-coupled live cell imaging and quantified the fate of preexisting Glt1 polymers upon acute removal of amino acids. Indeed, switching aged cells from rich SC media to amino acid-deprived media caused a systematic and progressive depolymerization of Glt1 polymers (**Fig. 3E-F**). Thus, Glt1 polymerization in aged cells is reversible.

### Glt1 polymerizes through self-association of symmetric complexes

To understand the nature of the Glt1 structures, we examined them using correlative light and transmission electron microscopy (CLEM). Importantly, the electron micrographs showed that the Glt1 polymers are composed of bundled filaments (**Fig. 3G**). This filamentous ultrastructure suggested that Glt1 assembly might occur at the polymer ends. To test this, we compiled a live-cell imaging assay to follow the fusion of two mating yeast cells carrying GFP- or mKate2-tagged versions of Glt1. By following the fate of distinct Glt1 polymers in the newly formed zygote, we found that the two polymers were able to join at their distal ends in a stable manner, providing evidence that the Glt1 polymers can grow at the tips through the addition of new subunits (**Fig. 3H**). To gain more insight into the polymerization dynamics, we measured polymer growth in individual cells over several hours (**Fig. 3I**). Fitting 76 polymer growth traces starting from their nucleation (distinguished as the first visible Glt1-GFP puncta) revealed a bi-phasic growth curve. In the initial 24-minute fast-growing state the doubling time of polymer elongation was 7.9 minutes, which was followed by a 22-times slower growth phase (**Fig. 3J**). Together, these data indicate that Glt1 undergoes a rapid switch-like transition from diffuse to filamentous polymers during the early aging of budding yeast cells.

To understand the structural basis of Glt1 polymerization, we modelled the yeast Glt1 structure based on the structure of the homologous *Azospirillum brasilense* GltS, which consists of two subunits that assemble into a hexameric oligomer with threefold dihedral (D3) symmetry ^57^. We found that the three LiP-MS identified Glt1 peptides that were protected from proteolytic cleavage in aged cells localized on the three-fold symmetric face of the GltS structure (**Fig. 3K**). We hypothesized that this face represents the polymerization interface mediating Glt1 self-assembly and becomes protected from proteolytic cleavage when Glt1 is polymerized. To test this hypothesis, we performed site-directed mutagenesis focusing on the LiP-MS identified region (LiP 1). Surface hydrophobicity can mediate the self-assembly of symmetric protein complexes ^58^, and we found that exchanging surface exposed hydrophobic residues for polar residues compromised Glt1 polymerization during aging (**Fig. 3L-M, Fig. S5C**). We focused on mutant 3 (H1537A, Y1538N, L1539S, henceforth *GLT1-MUT*) and validated that it displayed similar expression and catalytic activity as wild type Glt1 (**Fig. S5D-G**), demonstrating that the mutation does not prevent polymerization through adverse effects on Glt1 expression or folding. Overall, our data suggest that Glt1 filaments form during aging via the polymerization of hexameric (D3 symmetry) or trimeric (cyclic, C3 symmetry) Glt1 homomers (**Fig. 3N, Fig. S5H-I**).

### Inhibiting Glt1 polymerization delays aging, prevents intracellular amino acid accumulation and attenuates mitochondrial dysfunction

Several metabolic enzymes undergo reversible polymerization as an adaptive response to changes in the environment ^59,60^. We considered whether age-related Glt1 polymerization is an adaptation that impacts cellular longevity. To test this, we compared the replicative life span of wild type cells to that of CRISPR-Cas9-edited cells expressing non-polymerizing *GLT1-MUT*. Interestingly, microdissection mediated life span analysis revealed that the polymerization-deficient *GLT1-MUT* cells displayed a 20 % extension in median life span as compared to the wild type cells (p < 0.05, **Fig. 4A**). These data suggest that the Glt1 transition from the diffuse to polymerized form may promote aging in yeast cells.

To understand the role of Glt1 polymerization in aging and overall cellular function, we first compared the transcriptome of young and aged wild type and *GLT1-MUT* cells using RNA sequencing (**Fig. 4B**). We observed 814 genes with altered expression in aged wild-type versus *GLT1-MUT* cells, whereas the expression of only 11 genes was altered between young cells where Glt1 is soluble (**Fig. S6A**). Gene set enrichment analysis of the 814 genes showed significant differences in metabolic processes, with amino acid metabolism representing the second most significant GO-category (**Fig. S6B**). Together with the sensitivity of Glt1 polymerization to amino acid levels, this suggested that the Glt1 polymerization status is coupled to amino acid metabolism. We explored this further via targeted LC-MS metabolic analyses of young and aged wild type and *GLT1-MUT* cells (**Fig. 4B**). Principal component analysis (PCA) of the steady-state metabolome showed that aged wild-type cells clustered away from young wild-type cells, indicating that aging correlates with changes in cellular metabolism (**Fig. 4C**). Interestingly, aged wild-type cells carrying Glt1 polymers were the most distal cluster from the young cells and segregated away from the aged *GLT1-MUT* cells, indicating that aged yeast cells with Glt1 polymers have a distinct metabolic phenotype. Notably, young wild-type and *GLT1-MUT* samples co-clustered (**Fig. 4C**), which, together with the RNA seq data (**Fig. S6A**), further demonstrate that the H1537A, Y1538N, L1539S substitutions affect polymerization but do not alter the function of soluble Glt1.

We examined metabolites by performing an orthogonal partial least square (OPLS) analysis and observed increased amino acid levels and reduced mitochondrial TCA cycle metabolites in aged wild-type cells relative to aged *GLT1-MUT* cells (**Fig. 4D-E**). Indeed, the steady-state levels of amino acids increased up to 7-fold during aging in wild-type cells but not in *GLT1-MUT* cells (**Fig. 4F**). We confirmed that these differences cannot be explained by changes in cell size between the wild type and mutant cells (**Fig. S7A**). It was recently shown that loss of vacuolar acidification leads to overflow of cytosolic amino acids in aged yeast cells ^22,61^. We found that 8 out of 14 of vacuolar V-ATPase subunits displayed a structural change during aging, in line with a potential change in its function during aging. Thus, we tested if Glt1 polymerization is associated with vacuolar decline. However, Glt1 polymerization status did not affect vacuolar pH (**Fig. S7B**). Furthermore, inhibiting vacuolar ATPase with concanamycin A (ConcA) did not induce Glt1 polymerization (**Fig. S7C**). We conclude that Glt1 polymerization-associated increase in cytoplasmic amino acids is not associated with vacuolar decline.

To understand the origin of the increased amino acids, we performed a tracing analysis, growing cells with ^13^C_5_-^15^N_2_ isotope-labelled glutamine for 30 min **(Fig. 4B)**. Glt1 catalyzes the transfer of one amino group from a glutamine molecule to alpha-ketoglutarate, generating two glutamate molecules while NADH is oxidized in the process (**Fig. 4G**). We thus quantified the fraction of labelled glutamate derived from the Glt1 reaction relative to the rest of the labelled glutamate species derived from the Gdh1 and Gdh3 reactions. We found no differences in labelled glutamate levels between young and aged wild type or *GLT1-MUT* cells (**Fig. 4H, Fig. S7E-F**), suggesting that Glt1 retains its catalytic activity when it is polymerized. However, we observed that glutamine uptake was significantly increased in aged wild-type cells that carried Glt1 polymers (**Fig. 4I, Fig. S7E, G**).

These results suggested that the increase in amino acid levels may be mediated via differential uptake of glutamine and possibly other amino acids. Indeed, our RNA-seq identified amino acid transporters as a significantly altered GO-cluster between aged wild-type and *GLT1-MUT* cells (p adj.< 0.05) (**Fig. S6B**). To test whether increased glutamine import is linked to increased expression of glutamine transporters, we measured the expression of broad-specificity amino acid transporters Gnp1, Dip5 and Gap1 in young and aged wild-type and *GLT1-MUT* cells. Interestingly, Gnp1 and Dip5 displayed approximately 2-fold higher protein expression in the aged wild-type as compared to *GLT1-MUT* cells whereas no changes were detected in Gap1 (**Fig. S8A**). Gnp1 has high affinity for glutamine ^62^. In accordance with our metabolic analysis, we found that the expression of Gnp1 increased with age, and its levels in wild-type cells were significantly higher than in *GLT1-MUT* cells (**Fig. S8B-C**). We propose that Glt1 polymerization contributes to increased expression of amino acid transporters, including Gnp1, to promote the uptake of glutamine and likely other amino acids in aged cells (**Fig. 4J**).

Aged cells showed an accumulation of many amino acids (**Fig. 4F**). Since glutamine can serve as an amino group donor for the synthesis of other amino acids, we examined whether aged cells utilize excess glutamine to synthesize other amino acids **(Fig. S7D)**. Briefly, we used the tracing data to monitor the transfer of the labelled amino group derived from glutamine. Importantly, a comparison of ^15^N-labelled amino acids between wild-type and *GLT1-MUT* cells showed a significant increase of labelled phenylalanine in aged wild-type cells, but not in aged *GLT1-MUT* cells or young cells (**Fig. 4J**). A similar trend was observed in leucine/isoleucine (**Fig. 4K**). These data provide evidence that Glt1 polymerization-mediated glutamine uptake contributes to the accumulation of other amino acids, particularly phenylalanine. Altogether, these data indicate that Glt1 polymerization is associated to large scale metabolic and transcriptomic rearrangements during aging that lead to amino acid accumulation in aged cells and a shortened life span.

We therefore wanted to understand how the Glt1 polymerization-associated metabolic rearrangements in aged cells relate to the regulation of longevity. To first understand how Glt1 polymerization relates to aging (**Fig. 4A)**, we inspected mortality rates to determine at which timepoint *GLT1-MUT* cells begin to diverge from wild-type cells. We found that *GLT1-MUT* specifically impacted early mortality, in line with the timing of initial polymer formation (**Fig. S9A**).

Mitochondria function begins to decline during early aging in budding yeast ^21,22^. Importantly, this decline in mitochondrial function is associated with an overflow of cytosolic amino acids ^22,61,63^. Thus we measured mitochondrial membrane potential in young and aged cells as a proxy for mitochondrial fitness ^64^ (**Fig. S9B**), and found that mitochondrial membrane potential declined with age, as expected ^21,22^ (**Fig. 4L and Fig. S9C**). In contrast, the non-polymerizing *GLT1-MUT* cells maintained high mitochondrial membrane potential during the early phases of aging **(Fig. 4L and Fig. S9C**). Moreover, mitochondria were fragmented nearly twice as often in aged wild type cells (9.7 %) as compared to aged *GLT1-MUT* cells (5.1 %) (**Fig. S9D**). We did not detect significant differences in mitochondrial respiration by oxygraphy measurements at this point, probably since most cells are fermenting in these conditions (**Fig. S9E**). We considered an option where the decline in mitochondrial membrane potential could itself act as a signal to promote Glt1 polymerization. To test this, we reduced mitochondrial membrane potential by treating cells with carbonyl cyanide m-chlorophenyl hydrazone (CCCP), but this treatment had no effect on Glt1 polymerization (**Fig S9F**). The combined data suggest that polymerization does not depend on mitochondrial dysfunction, but that lack of polymerization can counteract mitochondrial dysfunction in aging cells.

Finally, we addressed the connection between Glt1 polymerization state and mitochondrial decline, testing the role of amino acid accumulation. We measured mitochondrial membrane potential in cells grown under moderate (SD media) and high (SD media + peptone) amino acid conditions that cause an elevation in intracellular amino acids, mimicking the effect of Glt1 polymerization in aged cells (**Fig. 4F**). Remarkably, culturing cells in high amino acid conditions reduced mitochondrial membrane potential in all cell populations, abolishing both age- and polymerization-associated differences (**Fig. 4M**). We infer that Glt1 polymerization compromises mitochondrial membrane potential via the elevation of intracellular amino acids.

## Discussion

Our findings demonstrate that mapping protein structural changes is a powerful way to identify mechanisms associated with cellular aging, which would have not been detectable with prevailing techniques that only quantify changes in transcriptome and proteome abundance. Since more than 80 percent of the structurally altered proteins have human homologs and showed enrichment for genes modulating life span in multicellular *C. elegan*s, the dataset has the potential to be broadly applicable.

The age-altered proteins identified in this study were enriched in essential, multifunctional proteins that occupied central positions in protein-protein interaction networks. Another interesting feature was the strong enrichment in proteins that can reversibly transition into condensates. Such proteins have emerged as major regulators of adaptive cellular responses that may adopt aberrant conformations during aging and age-related diseases ^11,12,65^. A subtype of condensates not previously linked to aging are structurally ordered polymeric assemblies made by metabolic enzymes. We show that surface-exposed hydrophobic residues mediate the self-assembly of symmetric Glt1 homomers during aging. The overall assembly mechanism of Glt1 is similar to that of several other homo-oligomeric proteins with internal symmetry ^58,66^, suggesting that cell intrinsic alterations in aging might increase the self-association potential of a broader range of proteins. It is not yet clear why aging leads to Glt1 polymerization. We provide evidence that the polymerization is reversible and sensitive to amino acids, perhaps reflecting an allosteric sensing mechanism. Additional cues affecting Glt1 polymerization may include potential changes in the intracellular pH ^55,67^.

We provide evidence that the Glt1 assemblies fulfill the criteria of being *bona fide* aging factors: 1. They form in response to aging, 2. They are asymmetrically segregated between aged mother cells and rejuvenating daughter cells, and 3. Preventing their formation extends life span ^15,39^. Our data show that Glt1 polymerization is associated with accumulation of amino acids suggesting that at least two separate pathways contribute to age-associated amino acid overflow: defective vacuolar compartmentalization ^22,61^ and Glt1 polymerization-mediated amino acid uptake and glutamine transamination. We found that amino acid accumulation is linked to mitochondrial dysfunction, potentially affecting the TCA cycle and mitochondrial iron-sulfur cluster proteins ^61,63^. Why then has polymerization been selected by evolution? In the wild, yeast cells do not grow in test tubes but in colonies formed by interactive communities where cells may engage in cross-generational amino acid exchange between young and aged cells ^68^. Thus, Glt1 polymerization could be an example of antagonistic pleiotropy that has evolved for the benefit of the community, but at the cost of aging of the individual cell.

In addition to affecting the mitochondria, we propose that age-related amino acid accumulation can lead to structural-functional alterations in proteins interacting with amino acids. For instance, 67 % of cytosolic aminoacyl-tRNA synthetases displayed structural changes during aging. This might reflect altered binding status to cognate amino acids, potentially affecting tRNA charging and fidelity of protein translation. Indeed, translation fidelity is important for life span regulation in eukaryotes ^31^ in part by overwhelming the proteostasis machinery ^32,51^. Our data support such a chain of events. We provide evidence that ribosome-associated Hsp70 chaperones involved in folding of nascent polypeptide chains are increasingly occupied by client interactions during aging. Similar changes were found in the ER Hsp70 Kar2/BiP and cytosolic Hsp70 Ssa1, likely reflecting increased occupancy of these chaperones by client proteins and their compartmentalization to aging cells ^25,47,69^. Such chaperone sequestering may lead to proteostasis collapse during later stages of aging ^47,70^.

Taken together, understanding the landscape of protein structural changes during aging provides opportunities to build novel hypotheses on how age-associated cellular changes at the pathway level with mechanistic precision.

### Limitations of the study

Our approach has some limitations that constrain the conclusions of our study. Our samples comparing two timepoints in aging limit the conclusions to early aging processes. Even though this is arguably the most informative way for capturing primary changes associated with aging, it will be important in the future to construct a more comprehensive temporal map of aging-associated protein structural changes. Although our data exposed known and novel processes related to aging, some of the detected changes can also derive from differences between the mother and daughter cells that are not related to aging *per se* ^71^. The proteome coverage in our LiP-MS analysis was 62.7 % of the expressed yeast proteome ^72^, so our study excludes a portion of the proteome, primarily consisting of low abundant proteins. Furthermore, owing partially to the limited resolution of the LiP-MS method, interpreting the nature of the structural change can be challenging, as they can be associated with various causes, for example altered interactions, folding or assembly change. Therefore, combining structural proteomics with secondary imaging-based approaches ^73^ could be a powerful approach, as shown also in this study for a limited set of proteins (Fig. S3). Finally, one intriguing challenge in the future is to gain a more comprehensive understanding of how Glt1 polymerization is regulated and how it inflicts transcriptomic and metabolic changes in aged cells.

## Key resources table

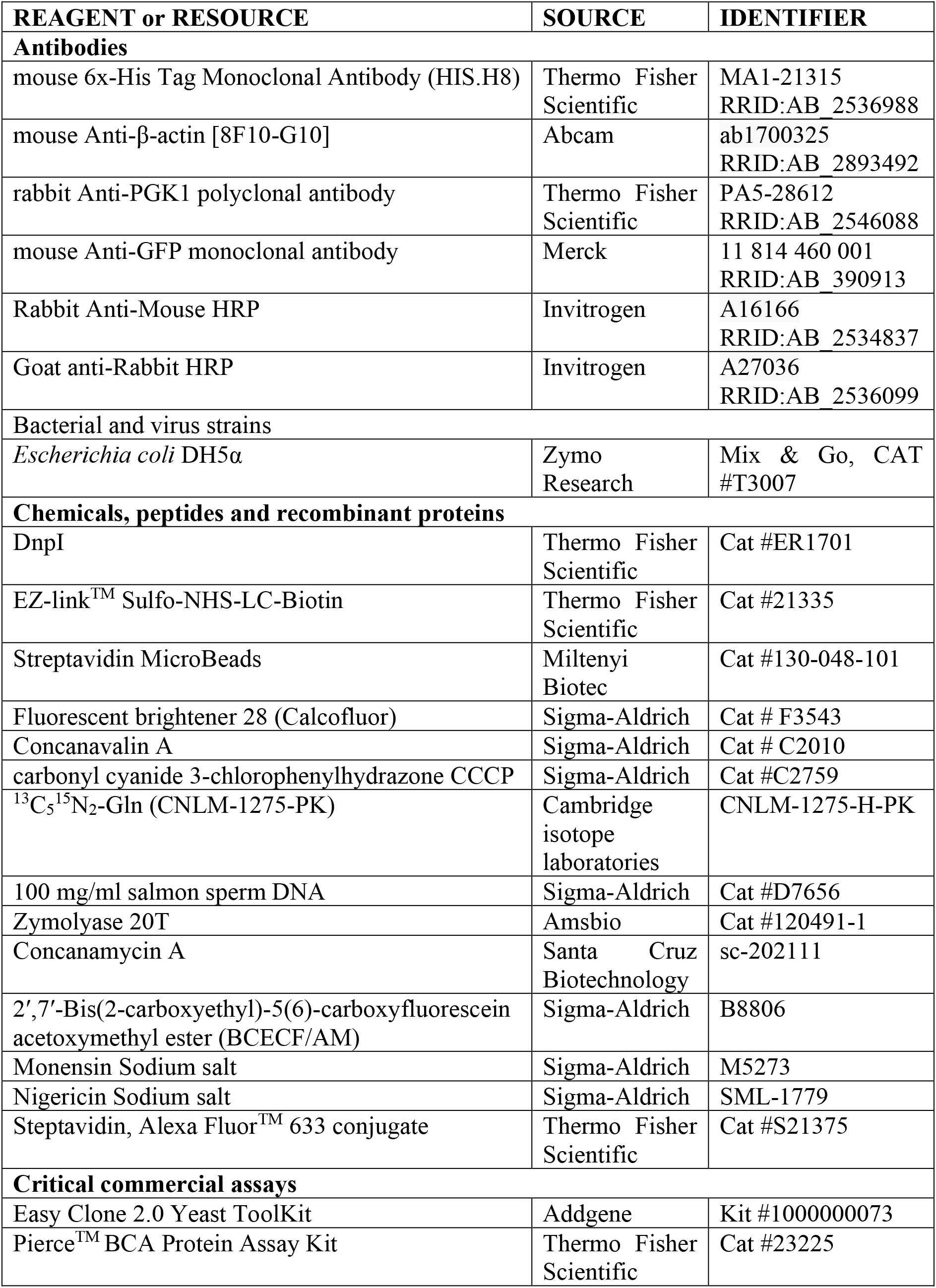
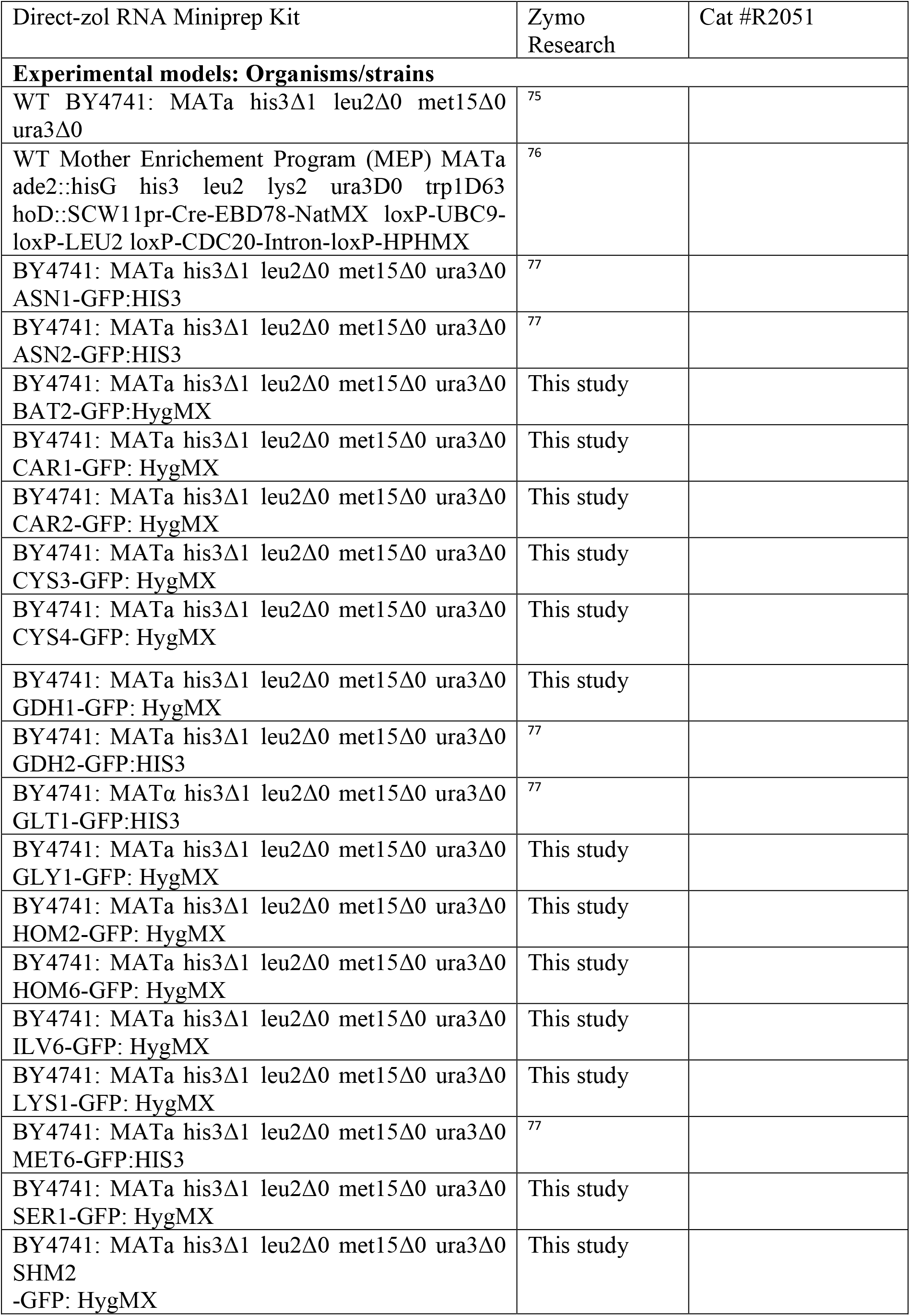
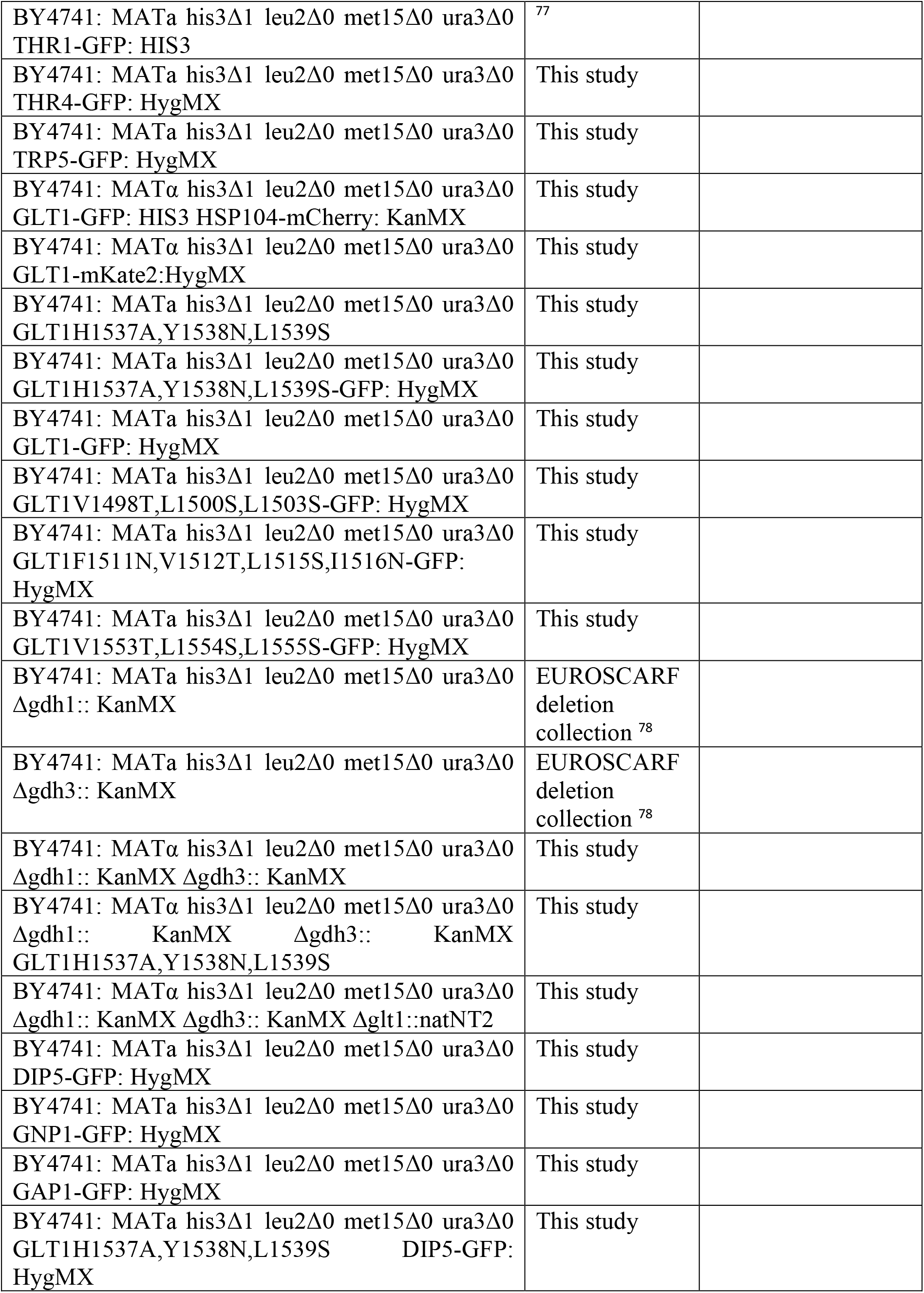
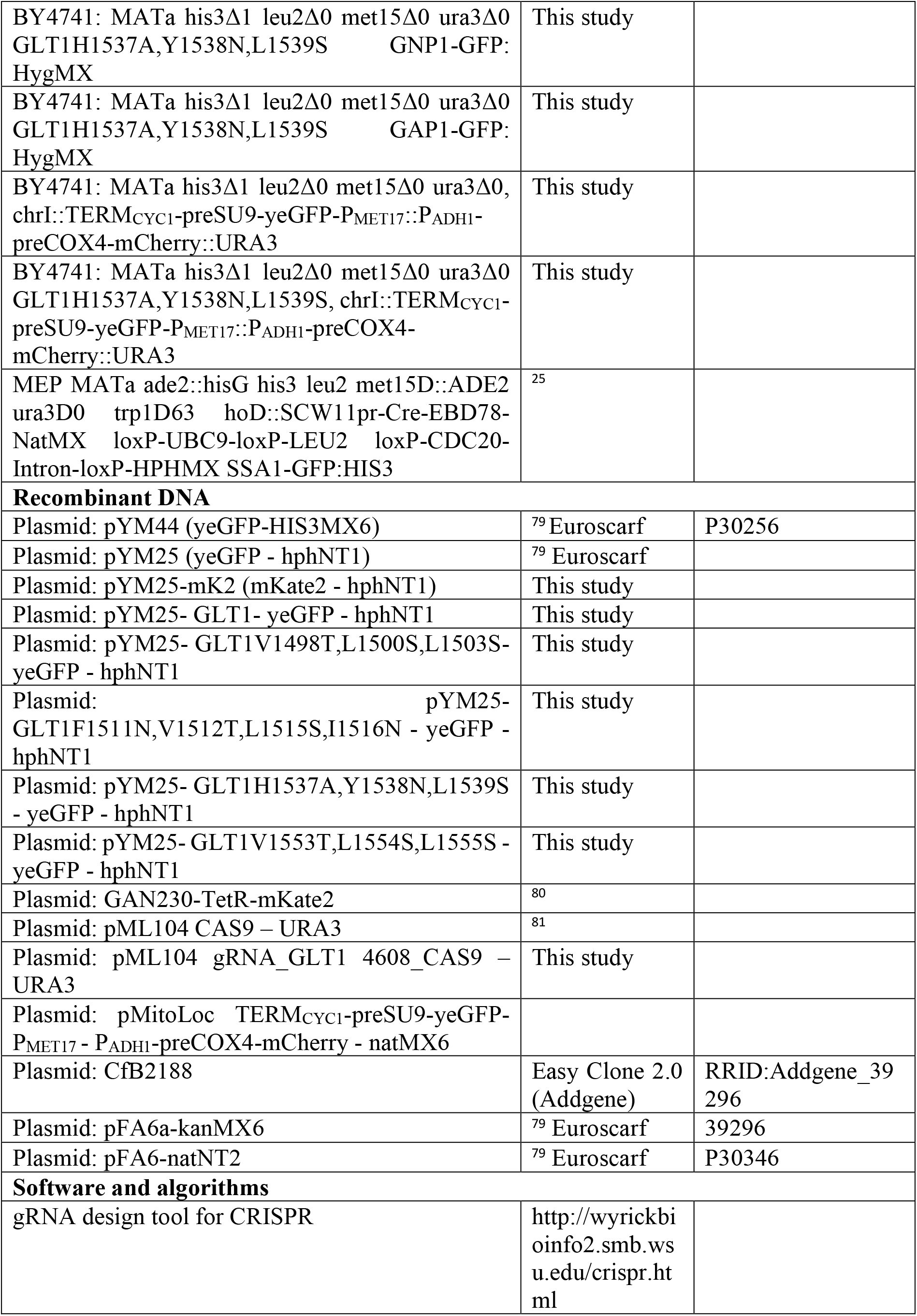
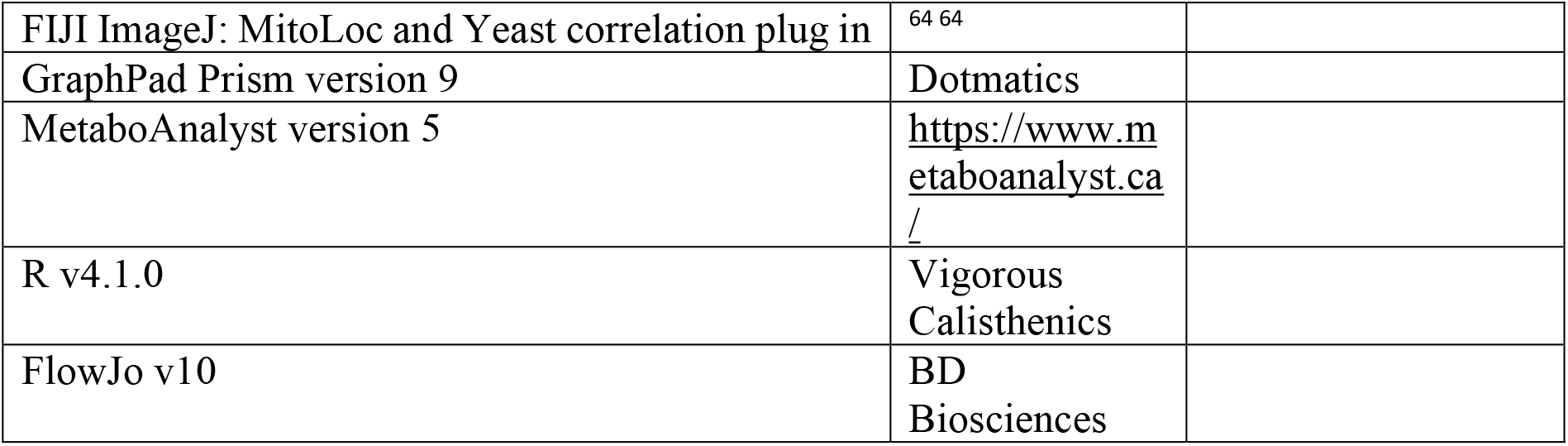

### Experimental model and subject details

#### Yeast strains and plasmids

All strains used in this study are derivatives of *Saccharomyces cerevisiae* S288c (BY4741) ^75^ and are listed in resource table. Endogenous gene tagging with GFP, mCherry or mKate2 were done by homologous recombination^79^. Briefly, primers annealing to the tagging cassette in the plasmid with additional 45 – 55 bp overhangs homologous to a place upstream and downstream of the STOP codon of desired gene were constructed. Tagging plasmid with mKate2 fluorophore YM25-mK2 was constructed by replacing GFP from YM25 plasmid^79^ with mKate2 from GAN230-TetR-mKate2 ^80^ plasmid by homologous recombination in *Escherichia coli* DH5α cells ^82^. For this, YM25 was linearized to exclude GFP by PCR and mKate2 was amplified from the plasmid GAN230-TetR-mKate2 with primers containing 25 bp overhangs complementary to linearized YM25. Purified fragments were assembled by mixing them vector:insert ratio of 1:3 and transforming them into *E. coli* DH5α strain ^82 82^. For genomic integration of MitoLoc, EasyClone 2.0 (Addgene) was used. Functional elements of original pMitoLoc TERM_CYC1_-preSU9-yeGFP-P_MET17_ - P_ADH1_-preCOX4-mCherry - natMX6 ^64^ was amplified with P1_f and G1_r primers and inserted in pCfB2188 plasmid. For genomic integrations into chromosome X, plasmid was cut with NotI and transformed into yeast (see Yeast transformation).

For Glt1 mutagenesis, GLT1 gene was inserted into GFP tagging plasmid YM25. First YM25 plasmid was linearized with PCR and assembled with GLT1 PCR fragment that contains 35 bp overhangs complementary to the plasmid by homologous recombination in *E. coli* DH5α strain ^82^. For this, purified PCR fragments were mixed with vector:insert ratio 1:3 and transformed to *E. coli* DH5α strain. Then primers with certain mismatches in GLT1 gene were designed so that the round-the-clock amplified YM25-GLT1-GFP plasmid would contain desired mutations. Reaction mixtures were treated with restriction enzyme DnpI (Thermo Fischer Scientific) to disrupt template DNA and transformed into *E. coli* DH5α strain. Newly generated plasmids were sequenced and inserted into yeast cells by homologous recombination with primers homologous to the sequence upstream and downstream of the gene (S1 and S2) to replace genomic GLT1 with GFP-tagged mutants (Janke *et al*., 2004).

Endogenous non-polymerizing GLT1 mutant H1537A, Y1538N, L1539S without tags or selection markers were generated with CRISPR-Cas9 system in yeast ^81^ and were used in experiments Fig. 4 and 5 and associated supplementary figures. Primers for gRNA were created with the online tool (http://wyrickbioinfo2.smb.wsu.edu/crispr.html) and the primer which was closest to the target sequence was selected at the position 4608 bp in the open reading frame. Vector for Glt1 mutation by CRISPR-Cas9 was prepared by ligating annealed and phosphorylated gRNA oligonucleotides with Smi/BclI digested plasmid pML104 ^81^. After sequencing, newly generated plasmid together with annealed 100 bp repair template containing desired mutation were transformed into yeast. Correct mutagenesis was confirmed with sequencing and Cas9 expressing plasmid was eliminated by streaking correct clone on YPD plate and selecting for single colonies which were no longer able to grow in absence of uracil.

#### Yeast cell culture, media and drug treatments

Cells were grown in exponential phase in YPD medium (1 % yeast extract, 2% Bacto peptone, 2% glucose) for 24 h before every experiment. Media were exchanged by washing cells twice with PBS and resuspending in the medium of interest. Media types refer to medium without amino acids (no AA) (6,7g/L YNB base, 2% glucose), minimal medium (MM) (0,02 g/L of Ura, Met, Leu, His, 6.7 g/L YNB base, 2% glucose), Synthetic minimal medium (SD) (0,02 g/L L-Ade, L-Arg, L-His, L-Met, L-Trp, Ura, 0.03 g/L L-Ile, L-Lys, L-Tyr, 0.05 g/L L-Phe, 0.1 g/L L-Leu, 0.15 g/L L-Val, 0.2 g/L L-Thr, 6.7g/L YNB base, 2% glucose), Synthetic complete medium (SC) (0.013 g/L L-Ade, 0.35 g/L L-Arg, L-inositol, 0.26 g/L L-Asp, L-Leu, 0.,057 g/L L-His, 0.52 g/L L-Ile, 0.09 g/L L-Lys, 0.19 g/L L-Met, 0.082 g/L L-Phe, L-Trp, 0.01 g/L L-Ser, 0.12 g/L L-Thr, L-Val, 0.013 g/L L-Ade, 0.018 g/L L-Tyr, 0.013 g/L L-Ade, 0.02 g/L Ura, 6.7 g/L YNB base, 2% glucose). Special additives were added to the final concentration of 2% Bacto peptone, 10 μM carbonyl cyanide 3-chlorophenylhydrazone (CCCP), 500 nM concanamycin A (ConcA).

## Methods details

### Transformation of yeast

Overnight yeast cell culture was diluted to OD_600_ 0.1 in liquid YPD media and grown until it reached OD_600_ 0.6. Then cells were pelleted by centrifuging them at 500 *g* for 5 min and washed twice with transformation buffer (100 mM LiOAc, 10 mM Tris, 1 mM EDTA, pH 8). Then cell pellet was resuspended in 72 μl transformation buffer and mixed with plasmid DNA (2 μl) or PCR product (10μl), 8 μl of 100 mg/ml salmon sperm DNA (prior to use boiled for 10min at 100 °C and cooled down on ice), 500 μl of PEG buffer (40% PEG-3350 (m/V), 100 mM LiOAc, 10 mM Tris, 1 mM EDTA, pH 8) and incubated in room temperature for 30 min. Then 65 μl of DMSO was added and yeast were subjected to heat shock at 42 °C for 15 min and spun down at 300 *g* for 5 min. If no auxotrophic marker was used for selection, yeast pellets were resuspended into 100 μl of corresponding media and plated on agar plates with the same media. If antibiotics were used, samples were incubated in 3 ml of YPD media for 2 – 3 h and then plated on YPD plates with antibiotics.

### Purification of young and aged yeast cells by magnetic-activated cell sorting

For young cell (0-1 generations) purification, cells were grown in exponential phase in YPD media for 24 h before biotinylation. Then 2 × 10^8^ cells were washed three times with PBS, resuspended in 3 mg/ml Sulfo-NHS-LC-Biotin (Thermo Fischer Scientific) solution in PBS and incubated on a rocking platform in room temperature for 30 min. Then cells were washed twice and resuspended in 200 ml of prewarmed YPD media. After 2.5 h of growth in 130 rpm at 30 °C, cells were harvested and resuspended into 2 × 10^8^ cells/ml in PBS with 2 mM EDTA and 1:20 of the volume of Streptavidin MicroBeads (Miltenyi Biotec). Cells were incubated on a rocking platform for 30 min in 4 °C and later resuspended into 12 ml of ice-cold PBS with 2 mM EDTA. Young cells were collected by passing cell suspension through the MS column (Miltenyi Biotec) attached to a magnetic stand and collecting flow through supernatant.

For aged cell (4-6 generations) purification, the procedure is the same, except that after biotinylation cells are grown in 500 ml of prewarmed YPD media for 6 h. Then when cells are put into the column, the column is washed 4 times with LiP buffer (20 mM hepes, 150 mM KCl, 10 mM MgCl_2_) for LiP-MS analysis and in PBS for metabolomics and RNA-seq analyses. Aged cells were eluted in LiP buffer or PBS by removing MS column from the magnetic stand.

After purification cells were resuspended into 1 ml of ice-cold LiP buffer or PBS and small amount of cell suspension was taken to access cell number with cell counter. The required number of cells was then divided into separate 1.5 ml tubes, spun down to remove supernatant and the pellet was snap frozen with liquid nitrogen for storage or directly used for downstream applications. In addition, a few microliters of initial purified cell culture were stained with calcofluor 10 μg/ml (from 1 mg/ml stock in DMSO) and fixed by adding two volumes of cell suspension of 4% PFA incubating cells for 15 min. In addition to calcofluor, streptavidin conjugated to Alexa Fluor™ 633 was added to the fixation samples for only aged cells for assessing the purity of purification by tracking initially biotinylated cells. After fixation, cells were centrifuged and the pellet was resuspended into 4 μl Mowiol-DABCO mounting media and put on a glass slide with thin glass cover slip. Cell age was assessed with microscopy (see Microscopy and live imaging) by calculating bud scars in calcofluor stained cell wall for at least 100 cells for each sample.

### Native protein extraction for LiP-MS

Cells were pelleted at 3000 *g* for 5 minutes at 4 °C and washed twice with ice-cold LiP-buffer (20 mM Hepes, 150 mM KCl, 10 mM MgCl_2_, pH 7.5), and subsequently resuspended in 1 mL LiP-buffer. Resuspended mixtures were snap-frozen in droplets by pipetting into liquid nitrogen. Lysis of the snap-frozen cell suspension droplets was performed using a 6775 Freezer/Mill Instrument (Thomas Scientific, Swedesboro, NJ, USA) according to the manufacturer’s instructions. Protein concentration of the resulted cell lysates (without clearing by centrifugation) were determined with bicinchoninic acid assay (BCA Protein Assay Kit, Thermo Scientific, Rockford, IL). Limited proteolysis (LiP) was performed as previously described^41^. Briefly, 100 µg of lysate was subjected to Proteinase K (Sigma Aldrich) treatment at an enzyme to substrate ratio (E:S) of 1:100 at RT. The LiP reaction was stopped by boiling in a water bath at > 95 °C. Complete protein denaturation was achieved by adding sodium deoxycholate (DOC) (Sigma Aldrich) to a final concentration of 5% (w/v). Disulfide bridges were reduced by adding tris(2-carboxyethyl)phosphine TCEP to a final concentration of 5 mM and incubation at 37 °C for 30 minutes. Reduced cysteine residues were alkylated by adding iodoacetamide IAA to a final concentration of 40 mM and incubation of 45 minutes at RT in the dark. Tryptic peptides were obtained by a first digestion with LysC (Wako Pure Chemical Industries) at an E:S of 1:100 for 4 h at 37 °C, followed by a second digestion with Trypsin (Promega) at an E:S of 1:100 overnight at 37 °C. For the digestion steps, DOC concentration and pH were adjusted to the optimal values as previously reported ^41^. Digestion was stopped by adjusting the pH to < 3 and precipitated DOC was removed by centrifugation. Finally, resulted peptides were desalted using Sep-Pak column (Waters) according to manufacturer’s instructions.

Peptides were separated using an online EASY-nLC 1000 HPLC system (Thermo Fisher Scientific) operated with a 15 cm long in house packed reversed-phase analytical column (Reprosil Pur C18 Aq, Dr. Maisch, 1.9 mm) before being measured on a Q-Exactive Plus (QE+) mass spectrometer. A linear gradient from 5%–25% acetonitrile in 140 min at a flowrate of 300 nl/min was used to elute the peptides from the column. Precursor ion scans were measured at a resolution of 70,000 at 200 m/z and 20 MS/MS spectra were acquired after higher-energy collision induced dissociation (HCD) in the Orbitrap at a resolution of 17,500 at 200 m/z per scan. The ion count threshold was set at 1,00 to trigger MS/MS, with a dynamic exclusion of 25 s. Raw data were searched against the S. cerevisiae Uniprot database using SEQUEST embedded in the Proteome Discoverer software (both Thermo Fisher Scientific). Digestion enzyme was set to trypsin, allowing up to two missed cleavages, one non-tryptic terminus and no cleavages at KP (lysine-proline) and RP (arginine-proline) sites. Precursor and fragment mass tolerance was set at 10 ppm and 0.02 Da, respectively. Carbamidomethylation of cysteines (+57.021 Da) was set as static modification whereas oxidation (+15.995 Da) of methionine was set as dynamic modification. False discovery rate (FDR) was estimated by the Percolator (embedded in Proteome Discoverer) and the filtering threshold was set to 1%. Label-free quantitation was performed using the Progenesis-QI Software (Nonlinear Dynamics, Waters). Raw data files were imported directly into Progenesis for analysis. MS1 feature identification was achieved by importing the filtered search results (as described above) from Proteome Discoverer into Progenesis to map the corresponding peptides based on their m/z and retention times. Annotated peptides were then quantified using the areas under their extracted ion chromatograms.

### 3D structural analyses

LiP conformotypic peptides were mapped to available protein structures to analyze the overlap of these structural alterations and functionally relevant sites within the protein (e.g., substrate binding sites). The EMBOSS Needle tool (EMBL-EBI) was used for pairwise sequence alignment of yeast protein sequences with homologous proteins. Structural analyses and visualization were performed with UCSF ChimeraX-1.1. ^83,84^. *Escherichia coli* DnaK protein structures were used to visualize conformotypic peptides of the yeast Hsp70 proteins (Ssa1, Ssb1, Ssb2 and Kar2) in open conformation (Protein Data Bank (PDB) ID: PDB:4B9Q) ^85^, closed conformation (PDB:2KHO) ^74^ and the substrate binding domain (SBD) in complex with a substrate peptide (PDB:1DKX) ^49^. The yeast Trm1 conformotypic peptide was mapped to a *Pyrococcus horikoshii* Trm1-S-adenosyl-L-Methionine complex structure (PDB:2EJT) ^52^.

A model of the Glt1 structure was created by SWISS-MODEL homology modeling server ^86^ using a homologous structure, glutamate synthases from *Azospirillum Brasilense* (PDB: 2VDC), as a template ^57^. Two types of Glt1 oligomers, with either C3 or D3 symmetry, were created by aligning the homology model with the symmetry-related chains in the D3 symmetric complex of the template structure using the Matchmaker tool in UCSF ChimeraX ^84^. Figures of the oligomers highlighting the peptides identified by LiP / MS and mutated residues were created in the same software.

### Contact site analysis

Chord plots illustrating contact sites between LiP peptide region in age-altered proteins and their clients were created using information on non-covalent contacts from the Protein Contact Atlas or custom data and processed with a python script to create a matrix of contacts between a protein and a client peptide. If suitable data was not available in Protein Contact Atlas, the non-covalent bonds were calculated with the coordinates provided in the PDB structure. Chord plots illustrations on client peptide bound to Hsp70s were based on *E. coli* DnaK (PDB:1DKX) ^49^ and chord plots on Trm1 bound to S-adenosyl-L-Methionine was based on Trm1 from *P. horikoshii* (PDB:2EJT) ^52^. Distances between pairs of atoms were defined with Euclidean distance. The radii of the atoms in contact were reduced from the distance. The radii of protein atoms were defined as previously described ^87^ and for client peptide atoms, Van der Waals radii were used. The threshold for non-covalent contact was set to 4 Å ^50^. The matrixes were imported to R 4.1.0 and plots were created with circlize package v0.4.15 chordDiagram function.

### Microscopy and microfluidics

Before imaging experiments, cells were grown in exponential phase in YPD media for 24 h and then were washed twice with PBS and resuspended in colorless media (for details see Yeast cell culture and media). Cells in colorless media were grown on a spinning wheel for 4.5 – 6 h and 1.26 × 10^6^ cells were stained with 100 μg/ml calcofluor for 5 min. Cells were washed once and resuspended into 1 ml of media. A small amount 50 μl of cell suspension was added to a well in a 96 thin glass bottom imaging well-plate pre-coated with concanavalin A (Sigma Aldrich) and topped up with 200 μl of colorless media. Well coating was performed by adding 100 μl of 2 mg/ml concanavalin A solution in (PBS with 50 mM of MnSO_4_ and 50 mM CaCl_2_), incubating in room temperature for 30 min and washing wells once with 150 μl of PBS. Imaging was performed with customized Olympus IX-73 inverted widefield fluorescent microscope DeltaVision Ultra (GE Healthcare) equipped with Pco edge 4.2ge sCMOS camera CentOS 7 Linux operating system. Imaging was done using 60x or 100x oil objectives and, depending on the fluorophore properties, with Blue (ex. 390 nm – em. 435 nm), Green (ex. 475 nm – em. 525 nm), Red (ex. 575 nm – em. 625 nm) and Far Red (ex. 632 nm – em. 679 nm) filter settings.

Microfluidic experiments were performed with CellASIC^®^ ONIX2 microfluidic operating system with CellASIC^®^ ONIX2 Y04C microfluidics plates, using manufacturer’s instructions. The imaging was performed with DeltaVision microscope as described above.

### Correlative light and electron microscopy (CLEM)

Yeast expressing Glt1-GFP were grown overnight in exponential phase in SD liquid media before the fixation. Then, 200 mL of the culture containing around 2.5 × 10^6^ cells was added to a 1 mg/mL Poly-L-Lysine (Sigma Aldrich) pre-coated MatTek dish with a gridded cover slip. The culture was incubated for 10 min at room temperature to allow cell sedimentation before removing the SD media and adding 200 mL of a fixative solution (3 % PFA, 0.5 % glutaraldehyde, 0.2 M citrate-phosphate buffer, pH 5.5). After 1 h incubation, samples were washed three times with 0.2 M citrate-phosphate buffer (pH 5.5) by keeping the washing solution on the sample dish for 5 min and 2 ml of the same buffer was added to fixed cells. Then cells were imaged with widefield fluorescent microscope DeltaVision Ultra (GE Healthcare) using brightfield channel and 60x oil objective (see Microscopy and live imaging) and cells of interest were located on the MatTek grid. Next, samples were fixed for electron microscopy by incubating the cells in fixative solution (2% glutaraldehyde, 3% PFA, 0.2 M citrate-phosphate buffer, pH 5.5) for 30 min at room temperature. After washing with 0.2 M citrate-phosphate buffer and distilled water, the cells were post-fixed with 2% potassium permanganate for 1 h, on ice. Prior gradual infiltration into low viscosity epoxy (TAAB, Aldermaston, UK) the samples were subjected to dehydration by increasing concentrations of ethanol. After incubating 2 × 3 h with 100% epoxy, the block was polymerized at 60°C for 16 h. The block was removed from the cover slip and a pyramid was trimmed according to the finder grid pattern transferred to the block surface. Serial 60-nm-thick sections were cut with an ultramicrotome (Leica EM Ultracut UC6i or UC7, Leica Mikrosysteme GmbH, Austria) and collected on Pioloform coated, single slot grids. The sections were post-stained with uranyl acetate and lead citrate and examined using a Hitachi HT7800 transmission electron microscope (Hitachi High-Technologies, Tokyo, Japan) operated at 100 kV, and a Rio9 CMOS-camera (Gatan Inc., AMETEK, Pleasanton, CA).

### tRNA isolation and m^2,2^G quantification by UPLC/MS

TRNA was isolated from young and aged cells (see Purification of young and aged yeast cells by magnetic-activated cell sorting) as previously described ^88^ with the following modifications. Briefly, approx. 10_8_ of young or aged cells were resuspended in 250 µL of 0.9% NaCl and lysed in an equal volume of acidic phenol (pH 4.3) with 1/5 volume of 1-bromo-3-chloropropane (BCP) and glass beads. Cells were sheared by vortexing for 5 min at full speed, followed by centrifugation for 15 min at 10 000 *g*, 22 °C. The aqueous phase was then transferred to a new centrifugation tube and re-extracted with acidic phenol/BCP. The re-extracted aqueous phase was collected and the volume was adjusted to 10 ml with equilibration buffer EQ (10 mM Tris-HCl pH 6.3, 15% ethanol, 200 mM KCl), after which it was applied onto a pre-equilibrated (EQ buffer containing 0.15% Triton X-100) Nucleobond AX-100 column (Macherey-Nagel). The column was washed twice with wash buffer WB (10 mM Tris-HCl pH 6.3, 15% ethanol, 300 mM KCl). TRNA was eluted with 10 mL of elution buffer EB (10 mM Tris-HCl pH 6.3, 15% ethanol, 750 mM KCl) into 2.5 vol. of 99.6% ethanol. The tRNA was precipitated O/N at −20 °C and pelleted by centrifugation for 30 min at 10 000 *g*, 4 °C. Residual salt was removed by washing the pellet with 80% ethanol. Then, tRNA pellet was air-dried at RT and re-suspended in RNase/DNase-free water. Dephosphorylated monoribonucleosides for MS analysis were generated as previously described ^89^. For data normalization, cleaved monoribonucleosides were spiked with _15_N-labeled ribonucleosides, which served as an internal standard (25 ng _15_N-labelled ribonucleosides per 100 ng of sample). Samples were analyzed as described in ^90^ using a Waters Acquity® UPLC system attached to a Waters Synapt G2 HDMS mass spectrometer via an ESI ion source.

### Metabolite profiling

Before extraction young and old yeast cells were purified with magnetic sorter (see Purification of young and aged yeast cells by magnetic-activated cell sorting). 2.5 × 10^7^ cells were taken for steady state metabolomics. For flux analysis 2.5 × 10^7^ cells were treated in SC media with labeled 6.5 mM ^13^C_5_^15^N_2_-Gln (CNLM-1275-PK) for 30 min pulse. The yeast suspension was washed twice with 1xPBS, and metabolites were extracted with 500μl ice-cold buffer acetonitrile:ddH_2_O (80:20). Subsequently, the samples were sonicated 2 min and vortexed for 30 s for 3 rounds in total, centrifuged 16 000 *g*, 10 min at +4°C and the supernatant was taken to further analysis.

All samples were analyzed on Thermo Q Exactive Focus Quadrupole Orbitrap mass spectrometer coupled with a Thermo Dionex UltiMate 3000 HPLC system (Thermo Fisher Scientific, Inc.). The HPLC was equipped with a hydrophilic ZIC-pHILIC column (150 × 2.1 mm, 5 μm) with a ZIC-pHILIC guard column (20 × 2.1 mm, 5 μm, Merck Sequant). 5 μl of the samples were injected into the LC-MS after quality controls in randomized order having every 10^th^ sample as blank. The separation was achieved by applying a linear solvent gradient in decreasing organic solvent (80– 35%, 16 min) at 0.15 ml/min flow rate and 45°C column oven temperature. The mobile phases were following, aqueous 200 mmol/l ammonium bicarbonate solution (pH 9.3, adjusted with 25% ammonium hydroxide), 100% acetonitrile and 100% water. The amount of the ammonium bicarbonate solution was kept at 10% throughout the run resulting in steady 20 mmol/l concentration. Metabolites were analyzed using a MS equipped with a heated electrospray ionization (H-ESI) source using polarity switching and following setting: resolution of 70,000 at *m/z* of 200, the spray voltages: 3400 V for positive and 3000 V for negative mode, the sheath gas: 28 arbitrary units (AU), and the auxiliary gas: 8AU, the temperature of the vaporizer: 280°C, temperature of the ion transfer tube: 300 °C. Instrument control was conducted with the Xcalibur 4.1.31.9 software (Thermo Scientific). The peaks for metabolites were confirmed using commercial standards (Merck Cambridge Isotope Laboratories & Santa Cruz Biotechnology). The final peak integration was done with the TraceFinder 4.1 SP2 software (Thermo Scientific) and for further data analysis, the peak area data was exported as excel file.

### Transcriptomic profiling

After young and aged cell purification (see Purification of young and aged yeast cells by magnetic-activated cell sorting), pellets containing 2.5 × 10^7^ cells were resuspended with 800 μl of TRI reagent (Zymo research) and placed in 2 ml Touch Micro-Organism Lysing Mix tubes (Omni). In total, 4 replicates for each group were used. Then, cells were mechanically disrupted with Precellys 24 Tissue Homogenizer (Bertin Instruments) at 6000 rpm for 15s for a total of 4 cycles with 2 min break on ice between cycles. RNA was further extracted using Direct-zol RNA Miniprep Kit (Zymo Research) following manufacturer’s instructions. RNA concentration and purity was measured by NanoDrop, and RNA samples were stored at -80 °C before use.

RNA-seq library preparation and sequencing were performed by the Sequencing Unit of Institute of Molecular Medicine Finland FIMM Technology Centre at the University of Helsinki. Illumina TruSeq Stranded mRNA library preparation Kit (Illumina) and NextSeq 500 Mid Output Kit PE75 (120 M reads) (Illumina) were used following manufacturer’s instructions.

### Protein extraction and Western Blot

Cells were harvested by centrifugation and the pellet containing 6 – 8 × 10^7^ cells was resuspended in 250 μl Lysis Solution (0,25 M NaOH, 1% β-mercaptoethanol (v/v)) and 160 μl of 50% Trichloroacetic acid. After 10 min incubation on ice, cells were centrifuged at high speed (21 500 *g*) for 3 min and the pellet was subjected to 1 ml acetone. Samples were centrifuged again and the pellet was resuspended into sample buffer (120 mM Tris-HCl pH 6.8, 2% SDS) and heated in 95 °C for 5 min. Small amount of samples was used for quantification of protein concentration with BCA Protein assay Kit (Thermo Fisher Scientific) and 6x loading buffer (48% glycerol and 0,03% bromphenolblue) was added to the rest of the sample. For each sample 10 – 15 μg of protein was loaded on a SDS-PAGE gel. For Glt1 detection via Western blot primary mouse Anti-GFP (Sigma-Aldrich) or mouse Anti-6x-His Tag (HIS.H8) (Thermo Fisher Scientific) antibodies were used. As a loading control primary rabbit Anti-PGK1 or mouse Anti-β-actin [8F10-G10] antibodies were used. For detection horseradish peroxidase (HRP) conjugated Rabbit Anti-Mouse (Invitrogen) or Goat anti-Rabbit (Invitrogen) secondary antibodies were used.

### Flow cytometry

Cells were grown in exponential phase for 24 h before biotinylation to distinguish aged cell fraction. For that, cells were washed three times with PBS and resuspended in 2 mg/ml Sulfo-NHS-LC-Biotin (Thermo Fischer Scientific) solution in PBS and incubated on a rocking platform in room temperature for 30 min. Then cells were washed twice and resuspended in colorless SD media at the density of 5 × 10^4^ cells/ml. Cells were grown in 30 °C for 10 h to facilitate aging of biotinylated cells. Then, approx. 2 × 10^6^ cells were taken and stained with Steptavidin-Alexa Fluor™ 633 at the final concentration of 20 μg/ml for 5 min. Cells were washed with PBS and resuspended in 1 ml of colorless SD media in a Falcon flowcytometry tube.

For vacuolar pH measurements, cells were incubated with 50 μM of 2′,7′-Bis(2-carboxyethyl)- 5(6)-carboxyfluorescein acetoxymethyl ester (BCECF/AM) for 30 min at 30 °C just before staining with Steptavidin-Alexa Fluor™ 633. Calibration curve was made my resuspending BCECF/AM stained cells in calibration buffer (50 mM MES, 50 mM HEPES, 50 mM KCl, 50 mM NaCl, 0.2 M ammonium acetate, 10 mM sodium azide, 110 uM monensin and 15 uM nigericin) of different pH (5.5, 6.0, 6.5, 7.0, 7.5). Cells were analyzed with BD LSRFortessa Flow Cytometer at the Flow Cytometry Unit in the University of Helsinki. GFP fluorophore was exited with 488 nm laser and optical filter 530/30 (515-545 nm), BCECF/AM was excited with 405 nm laser optical filter 525/50 (500-550 nm) and 488 nm 530/30 (515-545 nm) and for Alexa Fluor™ 633 fluorophore 640 nm laser and 670/30 (655-685 nm) optical filter were used. Samples were collected by gating aged Steptavidin-Alexa Fluor™ 633 positive cells and collecting 10 000 events per sample. Flow cytometry data analysis was performed with FlowJo v10 software. Vacuolar pH of BCECF/AM stained cells was determined from signal in 488 nm /405 nm ratio and interpolated from the calibration curve using GraphPad v9 software.

### Replicative life span assay

Single cells of wt and non-polymerizing Glt1 mutant (H1537A, Y1538N, L1539S) were placed on YPD plate. After the first division, the daughters were selected for the aging experiment and the mother cells were discarded. These selected cells were then followed through their replicative lifespan by removing newly divided daughter cells. Cell replicative life span was followed for 12 h in 30 °C following by incubation in + 4 °C overnight. Finite number of divisions for each cell were then plotted to generate the survival curves. The experiment was conducted with 3 biological replicates, N_mutant_ = 62, N_wt_ = 40.

### Oxygen consumption assay

After young and aged cell purification (see Purification of young and aged yeast cells by magnetic-activated cell sorting) equal amount of 1.4 * 10^7^ cells were placed in 2 ml of SD media (see Yeast cell culture and media) and incubated with shaking at 30 °C for 30 min. Then 1 ml of cell culture was placed in Oxygraph+ system (Hansatech Instruments) and oxygen consumption was measured for 4 min from each strain. As a control 10 μM of mitochondrial respiration inhibitor carbonyl cyanide 3-chlorophenylhydrazone CCCP was added at the end of the measurement to account for false positive signal. In total two biological replicates were used for each strain.

## Quantification and statistical analyses

### LiP-MS

Pairwise comparisons were performed between two conditions (young and aged), for PK-treated and control (non-PK) samples respectively. Peptide (PK samples) and protein (control samples) fold changes were calculated using three biological replicates per condition where the statistical significance was assessed with a two-tailed heteroscedastic Student’s t test. A fold change was considered significant with a q-value (FDR-corrected p-value) < 0.05 for peptides and proteins ^91^. To correct for protein abundance changes, a normalization factor based on q-value filtered protein fold change for each protein was applied on corresponding peptide intensities. After correction, prefiltered peptides with an absolute abundance change > 5 were selected as candidates. Functional enrichment by overexpression analysis of significant LiP hits were done with clusterProfiler v4.0.0 in R 4.1.0.

### Analysis of characteristics enriched among age-altered proteins

Most parameters for characterization of age-altered proteins were extracted from previously collected datasets described in ^92^. Specifically, information about protein-protein interactions were obtained from BioGRID database ^93^, number of GO functions retrieved through GO Slim annotations ^92^, protein complex network using physical interactome map of yeast ^94^, mutation rate from whole genome sequencing ^95^, protein abundance were obtained from cells grown in log-phase and rich media from GFP signal of endogenously GFP-tagged proteins ^96^ and translation rate as a function of ribosome density and occupancy ^97^. The lists of anti-longevity genes of *C. elegans* and *S. cerevisiae* were obtained from GenAge database ^98^ and the yeast list was supplemented with genes identified in ^99^. Statistical significance for these comparisons was assessed by unpaired t-test with Welch correction with False Discovery Rate (FDR) for multiple discoveries. Effect size is displayed as common language effect size (CLES). Information about protein solubility were obtained from SILAC based MS measurements ^100^, aggregation during heat stress ^44^, stress granule formation from affinity purification ^101^, asymmetrically retained long-lived proteins and their fragments (LARPs) from stable-isotope pulse-chase and total proteome MS ^45^, essential genes from growth of deletion strains in rich medium ^102^ and proteins with PEST motifs were predicted using EMBOSS 6.5.7 ^92^. Age-altered and background proteins were categorized with these characteristics by constructing contingency tables. The sample size n for individual comparisons between age altered proteins (N = 468) and background proteins (N = 2365) varies depending on data availability. Statistical significance for contingency tables was assessed with Fisher’s exact test and effect size is displayed as odds ratio (OR). Outliers were determined with Robust regression and outlier removal (ROUT) test (Q = 1%).

### Microscopy image analyses

Imaging data was analyzed with FIJI ImageJ software. MitoLoc analysis was performed using Yeast correlation plugin ^64^. Statistical significance in various images was calculated using GraphPad software. Two tailed unpaired t-test was used to assess statistical significance between two different groups. Ordinary one-way ANOVA with Dunnett’s correction for multiple comparisons was used to assess statistical significance for number of Glt1 polymers in different conditions. Two-way or three-way ANOVA with Šidák’s correction for multiple comparisons was used to assess statistical significance of mitochondrial function in young and aged WT and *GLT1-MUT* in different conditions. Correlation coefficient (R) and its significance for mitochondrial function dependance on age was computed with standard Pearson’s correlation test.

### Metabolomics

Peak intensity values were analyzed with Metaboanalyst software. Data was log2 transformed and Pareto scaled for principal component (PCA) and orthogonal partial least square (OPLS) analyses. Statistical comparison of metabolite peak intensities between WT and *GLT1-MUT* was done with one-way ANOVA with Dunnett’s correction for multiple comparisons using GraphPad v9 software.

### RNA-seq data analysis

Quality control of raw reads were performed with FastQC v0.11.9, followed by read filtering with Trim Galore v0.6.6 and rRNA removal with SortMeRNA v4.2.0. Transcriptome mapping was done with Salmon v1.4.0 and quality control of read mapping was done with STAR v2.7.8a and Qualimap v2.2.2d. Summary of quality control results was reported with MultiQC v1.9. Differentially expressed genes between young and aged cells as well as between WT and *GLT1-MUT* cells were found using DESeq2 v1.32.0 (adjusted p-value < 0.05, Wald test) in R 4.1.0. Functional enrichment of differentially expressed genes by overrepresentation analysis was done with clusterProfiler v4.0.0. For this study, the yeast genome/transcriptome references and gene annotations from Ensembl release 103 were used, which are based on yeast S288C genome assembly R64-1-1.

## Data availability

The LiP-MS proteomics, transcriptomics and metabolomics data generated and analyzed in the this study are available in the Harvard Dataverse repository (https://dataverse.harvard.edu/privateurl.xhtml?token=d4325d32-19dc-473a-952a-359b2ad668cb). All other data are included within the Article and its Extended Data.

## Code availability

None of previously unreported custom code was generated in this study. Code used for processing of RNA sequencing data can be found in Harvard Dataverse depository (https://dataverse.harvard.edu/privateurl.xhtml?token=d4325d32-19dc-473a-952a-359b2ad668cb).

## Acknowledgments

We thank the members of the Saarikangas lab and Jette Lengefeld for input and Cory Dunn, Markus Ralser, Ahmad Khalil and Daniel Gottschling for reagents. We are grateful to the assisting personnel at the Light Microscopy (LMU), Electron Microscopy (EMBI) and Viikki Flow Cytometry FIMM Genomics and DNA sequencing and Genomics (BIDGEN) units and MIBS and BI Media Kitchens at the University of Helsinki. Graphics in Fig. 1A and Fig. 4B, Fig. S9B were made in ©BioRender - biorender.com. This work was supported by the University of Helsinki, the Academy of Finland (317038), Sigrid Jusélius Foundation, the Human Frontiers Science Program Young Investigator Grant (RGY0080/2020) and Doctoral Programme in Integrative Life Science.

## Author contributions

Conceptualization: JuS, YB, PP, JP. Methodology: JP, RMLC, YF, KT, DS, EE, PG, MD, MS, JeS, AL, JJ, HV, AN. Investigation: JP, RMLC, YF, KT, DS, EE, PG, MD, MS, AL, JH, JuS. Funding acquisition: JuS. Supervision: PT, LH, EJ, AK, PLS, VH, PP, YB, JuS. Writing - original draft: JP, JS with feedback from all the authors.

## Supplemental information

## Supplemental figure legends

**Figure S1. Age-related structural changes are prevalent in abundant and pleiotropic regulators of metabolism**.

**A**. Overview of the online ProtAge database containing all age-altered proteins found at https://protage-server-21.it.helsinki.fi/.

**B**. Number of link communities in protein-protein interaction (PPI) network between age-altered proteins (*n* = 460) (light blue) and unaltered proteins (*n* = 2279) (grey).

**C**. Fraction of essential genes among age-altered proteins (*n* = 468) (light blue) compared to unaltered proteins (*n* = 2332) (grey).

**D**. Mutation rates expressed as density of amino acid substitutions between age-altered protein coding genes (*n* = 417) (light blue) and unaltered (*n* = 2198) (grey).

**E**. Distribution of protein solubility between age-altered proteins (*n* = 383) (light blue) and unaltered proteins (*n* = 922) (grey).

**F**. Comparison of protein abundance expressed as copy numbers per cell between between age-altered proteins (light blue) (*n* = 361) and unaltered proteins (*n* = 1572) (grey).

**G**. Relative translation rate of age-altered proteins (*n* = 444) (light blue) and unaltered proteins (*n* = 2162) (grey).

**H**. Fraction of proteins with protein degradation PEST motif between age-altered proteins (*n* = 468) (light blue) and unaltered proteins (*n* = 2332) (grey). Black line in box plots in (B, D, F, G) is median and whiskers mark 10 and 90 percentile values. Effect size is displayed as common language effect size (CLES) (B, D, F, G), and as odds ratio (OR) (C, E, H). *\*P* < 0.05; *\*\*\*\*P* < 0.0001.

**Figure S2. Mapping aging-associated structural changes in the substrate-binding domain of Hsp70 chaperones**.

**A**. Interconnected network among age-altered regulators of protein folding GO category. The graph depicts high confidence protein-protein interactions (STRING, confidence score > 0.7).

**B**. Cartoon representation of ATP-bound ‘closed’ Hsp70. The close-up displays the Lip-peptides in Ssa1 (red) located in the nucleotide binding domain (NBD) (gray) (PDB:4B9Q; ^74^).

**C**. Ssb1 LiP-peptides (red) that localize in the substrate binding domain (SBD) of client peptide (yellow)-bound Hsp70 (PDB:1DKX) ^49^.

**D**. Chordplot illustration based on (PDB:1DKX) ^49^ of contacts between Ssb1 LiP-peptides (red) and client peptide (yellow). Contacting secondary structures are illustrated by inner arcs.

**E**. Ssb2 LiP-peptides (red) localizing in the substrate binding domain (SBD) of client peptide (yellow)-bound Hsp70 (PDB:1DKX) ^49^.

**F**. Chordplot illustration based on (PDB:1DKX) ^49^ of contacts between Ssb2 LiP-peptides (red) and client peptide (yellow). Contacting secondary structures are illustrated by inner arcs.

**G**. Kar2 LiP-peptides (red) localizing to substrate binding domain (SBD) of client peptide (yellow)-bound Hsp70 (PDB:1DKX) ^49^.

**H**. Chordplot illustration based on (PDB:1DKX) ^49^ of contacts between Kar2 LiP-peptides (red) and client peptide (yellow). Contacting secondary structures are illustrated by inner arcs.

**Figure S3. Age-related structural change maps to conserved methyl donor-binding site of Trm1**.

**A**. Protein sequence alignment between *Pyrococcus horikoshii, Saccharomyces cerevisiae* and *Homo sapiens* shows high conservation in the Motif II region where LiP-peptide is located. Stars mark critical residues (Ile79 and Asp78) for methyl donor binding.

**B**. Illustration of Trm1 mediated N2,N2-dimethylguanosine (m^2^,^2^G) modification at a position G_26_ in tRNAs.

**Figure S4. Microscopic evaluation of amino acid synthesizing enzymes identifies age-related polymerization of glutamate synthase Glt1**.

**A**. Enrichment map of overrepresentation of age-altered proteins in amino acid metabolism. The size of dots represents the number of proteins for every group. The color change from blue to red signify the size of an adjusted p value (p.adj. < 0.05).

**B**. GFP-tagged amino acid metabolizing enzymes (green) that appeared soluble and showed no visible localization changes with age. Cell age in (B-D) can be determined from bud scars on cell wall stained with calcofluor (magenta). Scale bar 1 μm.

**C**. GFP-tagged amino acid metabolizing enzymes (green) that showed punctate appearance. Scale bar 1 μm.

**D**. Quantification of Glt1 polymers in aged cells (4 – 6 divisions) when tagged with either GFP or mKate2 (monomeric) fluorophores. Number of polymers displayed as mean ± SD. *n*_*(GFP)*_ = 82, *n*_*(mKate2)*_ = 36.

**E**. Representative image of aged cells expressing Glt1-GFP (green) and Hsp104-mCherry (red) show no colocalization between Glt1 polymers (arrowhead) and age-associated aggregates. 52 cells were examined, and none displayed accumulation of Hsp104-mCherry in Glt1 polymers. Scale bar 2 μm.

**Figure S5. Surface exposed hydrophobic residues mediate Glt1 self-assembly**.

**A**. Quantification of cells with Glt1 polymers in normal (2% glucose (glc)) and glucose limitation (0.1% glc) conditions in synthetic complete media. Error bars represent mean ± SD from 3 biological replicates. *n* = 452 for each group.

**B**. Quatification of cells with Glt1 polymers in SD medium containing 5 g/L of (NH_4_)_2_SO_4_ as high nitrogen (high NH_4_) conditions or SD medium with twice less of yeast nitrogen base (YNB) as low nitrogen (low NH_4_) conditions. *n*_*SD*_ = 558, *n*_*High NH4*_ = 164, *n*_*Low NH4*_ = 214 from three biological replicates.

**C**. Validation of the effect of mutagenesis on Glt1 self-assembly. Representative images of Glt1 mutant cells. Calcofluor marks the cell wall (magenta). Scale bar 1 μm.

**D**. Western blot analysis of expression levels of Glt1 mutant proteins. Glt1-GFP was detected with anti-GFP antibody (band size 265 kDa). Antibody against Pgk1 (45 kDa) was used as a loading control.

**E**. Quantification of relative band intensity of Glt1 mutant M3 (*GLT1-MUT*) to the WT. Bands were normalized to the band intensity of loading control. *n* = 2.

**F**. Schematic overview of L-glutamate synthesis in yeast which relies on two pathways: glutamate synthase Glt1 and glutamate dehydrogenases Gdh1 and Gdh3.

**G**. Catalytic activity of Glt1 mutant M3 (*GLT1-MUT*) was evaluated by growth essay on media without glutamate in strains that were deleted of *GDH1* and *GDH3*. Glt1 mutant M3 (*GLT1-MUT*) rescues the growth defect observed in a triple deletion (∆g*dh1* ∆*gdh3* ∆*glt1*) in a comparable manner as WT Glt1.

**H**. Localization of LiP-peptides in the cartoon model of Glt1 hexamer with dihedral symmetry (D3) and putative assembly mechanism.

**I**. Localization of LiP-peptides in the cartoon model of Glt1 trimer with cyclic symmetry (C3) and putative assembly mechanism. Error bars represent mean ± SD. Significance of the difference in A-B, E) was assessed with unpaired two-tailed t test assuming equal variance. n.s. not significant.

**Figure S6. Glt1 polymerization mediates transcriptional changes in aged cells**.

**A**. Table summarizing number of genes with significantly age-altered expression levels between WT and *GLT1-MUT* cells Significance of differential expression was statistically evaluated with Wald test. (*P* adjusted < 0.05), *n* = 4.

**B**. GO enrichment by over-representation analysis between aged WT and *GLT1-MUT* cells showing enriched biological process in which gene expression was altered when polymerization of Glt1 is engaged. The size of dots represents the number of proteins for every biological process. The color change from blue to red signify the size of an adjusted p value (*P* adjusted < 0.05).

**Figure S7. Glt1 polymerization results in increased glutamine uptake in aged WT cells**.

**A**. Cell size distribution among aged cells with or without their budding daughters between WT and *GLT1-MUT*. Number of cells analyzed: aged mother cells *n* = 84, aged mother cells with their budding daughters *n*_*WT*_ = 138 *n*_*GLT1-MUT*_ = 130.

**B**. Vacuolar pH in WT and *GLT1-MUT* cells in the presence or absence of 500 nM of concA measured by flow cytometry in cells stained with 50 μM of 2′,7′-Bis(2-carboxyethyl)-5(6)-carboxyfluorescein acetoxymethyl ester (BCECF/AM). Biological replicates *n* = 3, analyzed cells per sample n ≥ 10000.

**C**. Quantification of cells with Glt1 polymers in the presence or absence of 500 nM concanamycin A (concA). Error bars represent mean ± SD from 3 biological replicates. *n*_*SD*_ = 339, *n*_*SD concA*_ = 366.

**D**. Schematic overview of labelled ^13^C_5_-^15^N_2_-glutamine incorporation into cell metabolism. Labelled glutamine (Gln) taken up by cells is used for glutamate (Glu) production by Glt1 and Gdh1 and Gdh3. Glutamate can be further used for glutamine and alpha-ketoglutarate (α-KG) synthesis or for amino acid (AA) synthesis via transamination.

**E**. Fractional contribution of labelled glutamate species from Glt1 or Gdh1 and Gdh3 reactions in young and aged cells. Squares represent parts of whole of total labelled glutamate species. *n* = 4.

**F**. Fractional enrichment of all glutamate species in WT and *GLT1-MUT* in young and aged cells. *n* = 4.

**G**. Fractional enrichment of all glutamine species in WT and *GLT1-MUT* in young and aged cells. *n* = 4. Error bars represent mean ± SD. Statistical significance in (A, C) was assessed by two-tailer unpaired t test assuming equal variance and (F-G) with ordinary one-way ANOVA with Dunnett’s correction for multiple comparisons and B with two-way ANOVA Šidák’s correction. *\*P* < 0.05; *\*\*P* < 0.01; n.s. not significant.

**Figure S8. Polymerization of Glt1 results in expression changes in general amino acid transporters**.

**A**. General amino acid transporters that import glutamine Gnp1, Dip5 and Gap1 were tagged with GFP and their expression levels in young and aged cells were measured with flow cytometry. Heatmap summarizes fold change with age in WT and *GLT1-MUT* cells. Biological replicates *n* = 3, analyzed cells per sample *n* ≥ 10 000.

**B**. Microscopic images of Gnp1-GFP (green) in aged WT cells and *GLT1-MUT* cells. Cell age is indicated by calcofluor staining of the cell wall (magenta). Scale bar 1 μm.

**C**. Quantification of GFP signal from Gnp1-GFP transporter by flow cytometry between WT and *GLT1-MUT* in young and aged cells. Error bars represent mean ± SD. Statistical significance was assessed with two-way ANOVA with Šidák’s correction for multiple comparisons. Biological replicates *n* = 3, analyzed cells per sample *n* ≥ 10 000. \*\*\*\**P* < 0.0001.

**Figure S9. Glt1 polymerization is associated with mitochondrial dysfunction at early stage in replicative lifespan**.

**A**. Probability of death comparison between WT and *GLT1-MUT* cells during the first 20 divisions. *n*_*wt*_ = 40, *n*_*GLT1-MUT*_ = 62.

**B**. MitoLoc enables single-cell analysis of mitochondrial function through colocalization of mitochondrial preSU9-GFP (green) and cytosolic preCOX4-mCherry (red) signals. The import of preCOX4, but not preSU9, depends on mitochondrial membrane potential.

**C**. Mitochondrial function between WT and GLT1-MUT cells was evaluated with MitoLoc construct and is displayed as a Pearson’s correlation coefficient of colocalization between mitochondrial peSU9-GFP and cytosolic preCOX4-mCherry. Non-polymerizing mutant maintain stable mitochondrial function during the early stage of replicative life span (4-6 divisions). *n*_*WT*_ = 211, *n*_*GLT1-MUT*_ = 184.

**D**. Distribution of mitochondrial morphology in young and aged WT and Glt1 mutant cells. Pie charts represent parts of whole of mitochondrial morphology distribution in group of cells. *n*_*WT*_ = 211, *n*_*GLT1-MUT*_ = 184.

**E**. Oxygen consumption in young and aged cells between WT and GLT1-MUT cells. Oxygen consumption was measured with oxygraph in exponentially grown culture after MACS purification of young and aged cells. *n* = 3 biological replicates.

**F**. Quantification of cells with Glt1 polymers in the presence or absence of 10 μM carbonyl cyanide m-chlorophenylhydrazone (CCCP), which disrupts mitochondrial membrane potential. 3 biological replicates with *n* = 100 for each group. Error bars represent mean ± SD. Statistical significance (C, E) was assessed with two-way ANOVA with Šidák’s correction for multiple comparisons. *\*\*P* < 0.01; *\*\*\*P* < 0.001 *\*\*\*\*P* < 0.0001; n.s. not significant.

**Movie S1**.

Time-lapse fluorescence microscopy of dividing yeast cells expressing endogenous Glt1 tagged with GFP (green). Newly born daughter cells are born with diffuse Glt1. After a few divisions, Glt1 transitions into polymeric assembly which is asymmetrically retained in the mother cells during cell division. Images were taken every 15 min for a total period of 315 min.

