## Supplementary figures and images for "Global analysis of aging-related protein structural changes uncovers enzyme polymerization-based control of longevity"

### Supplemental figures S1 to S9

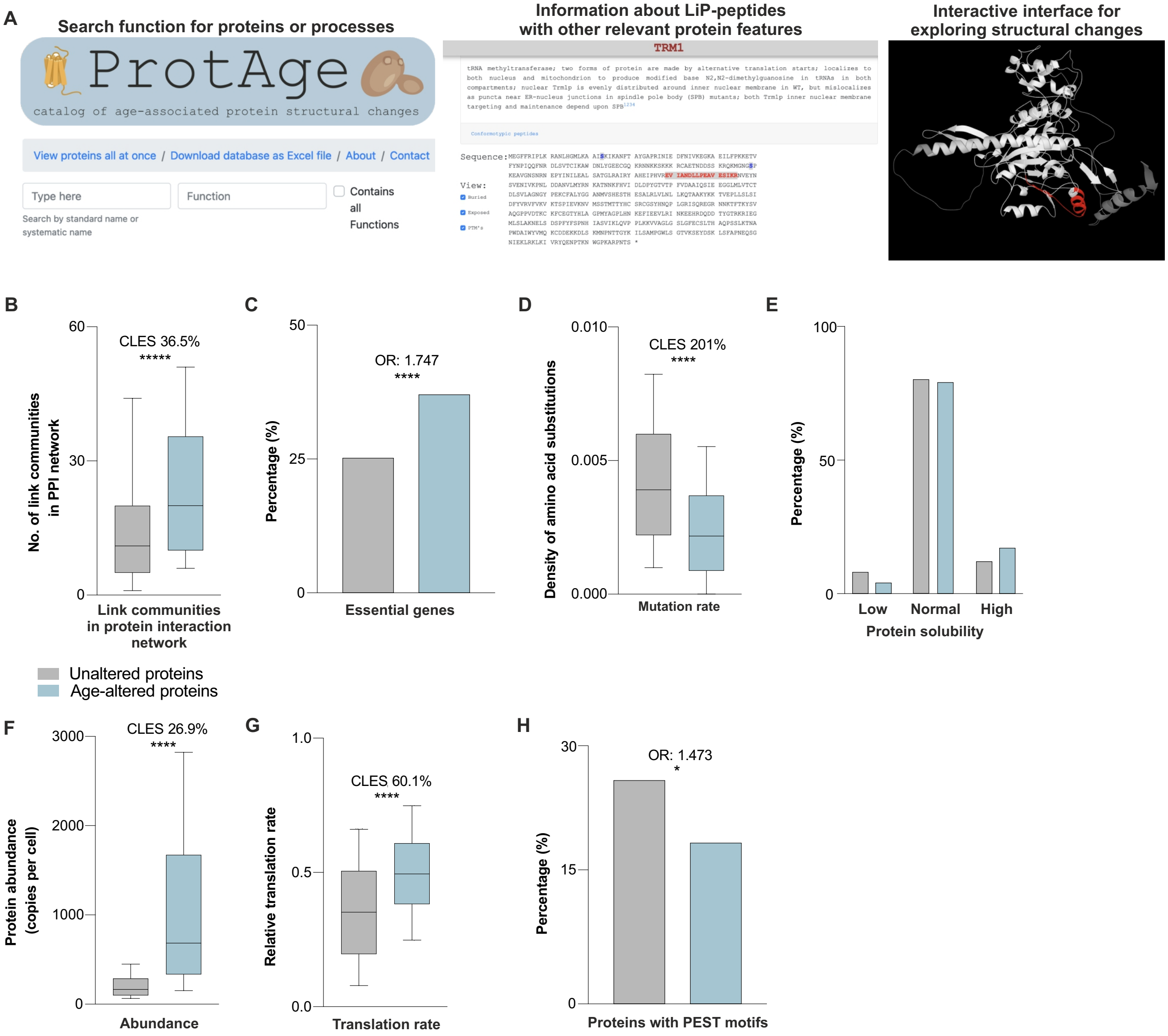

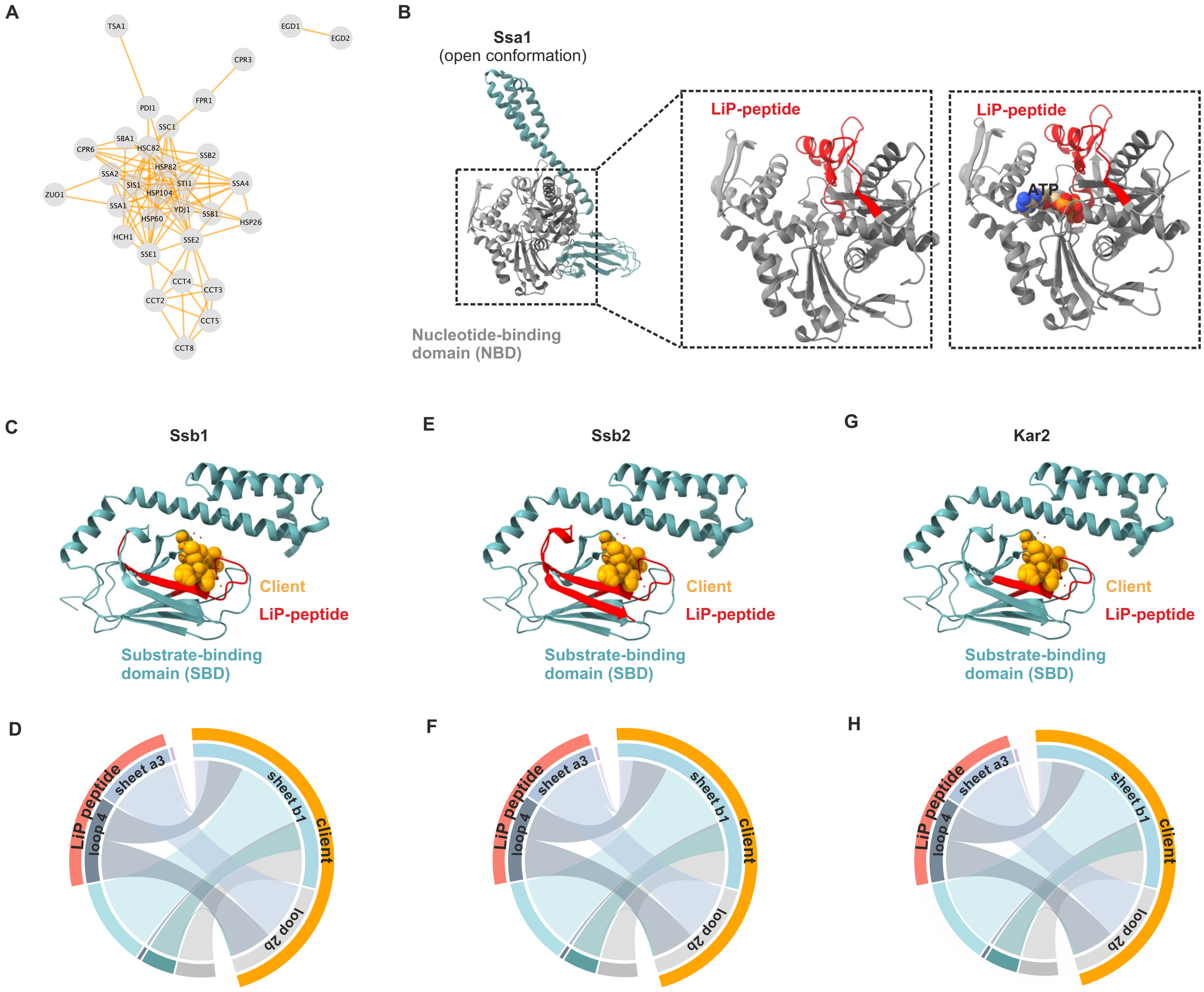

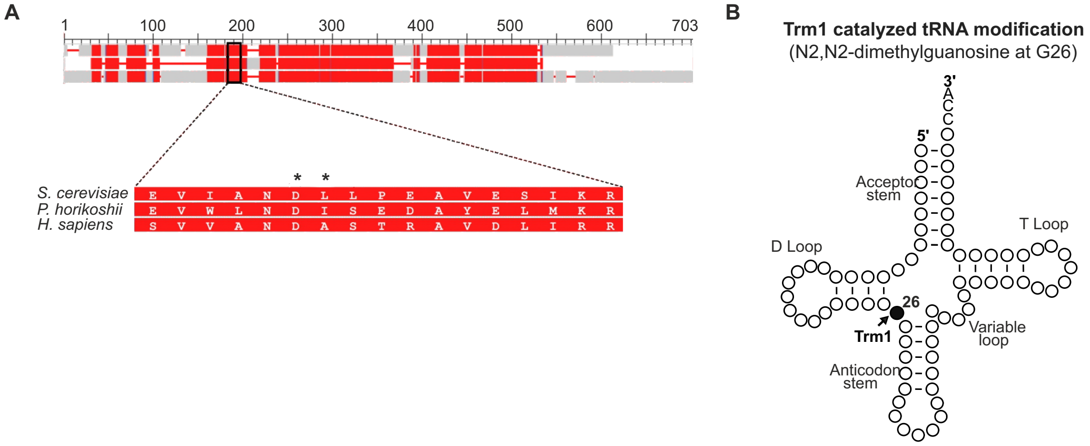

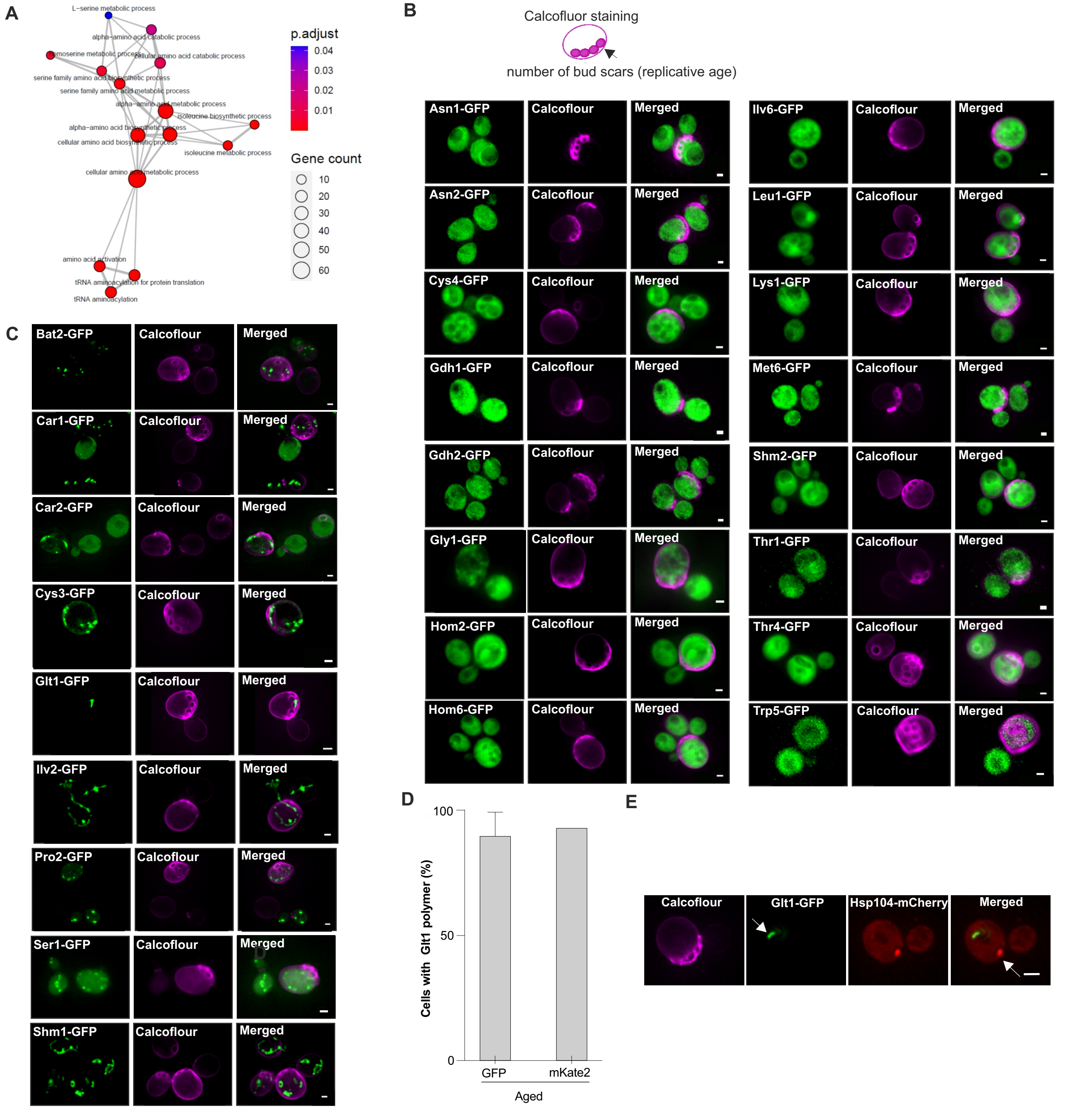

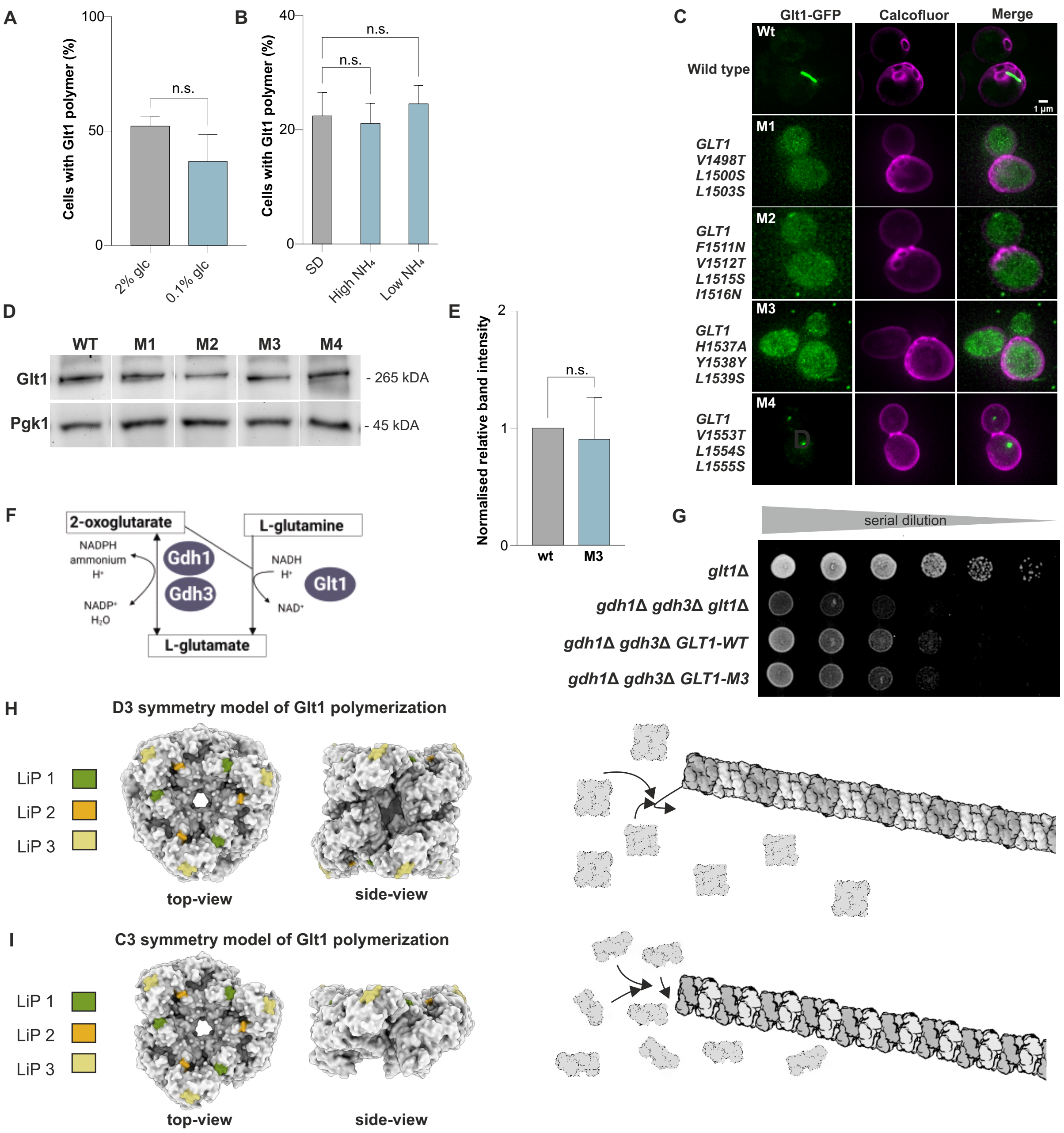

**A**

| **Sig . genes** | **WT vs.** | ***GLT1-MUT*** | **Old vs. Young** | |
|---|---|---|---|---|
| | Young | Old | wt | *GLT1-MUT* |
| Up | 9 | 337 | 471 | 543 |
| Down | 2 | 477 | 630 | 650 |
| Total | 11 | 814 | 1101 | 1193 |

**B**

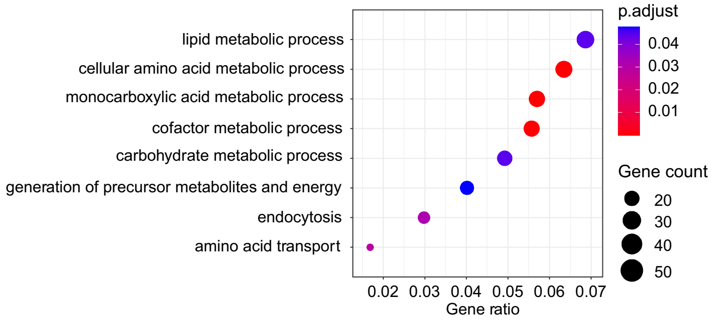

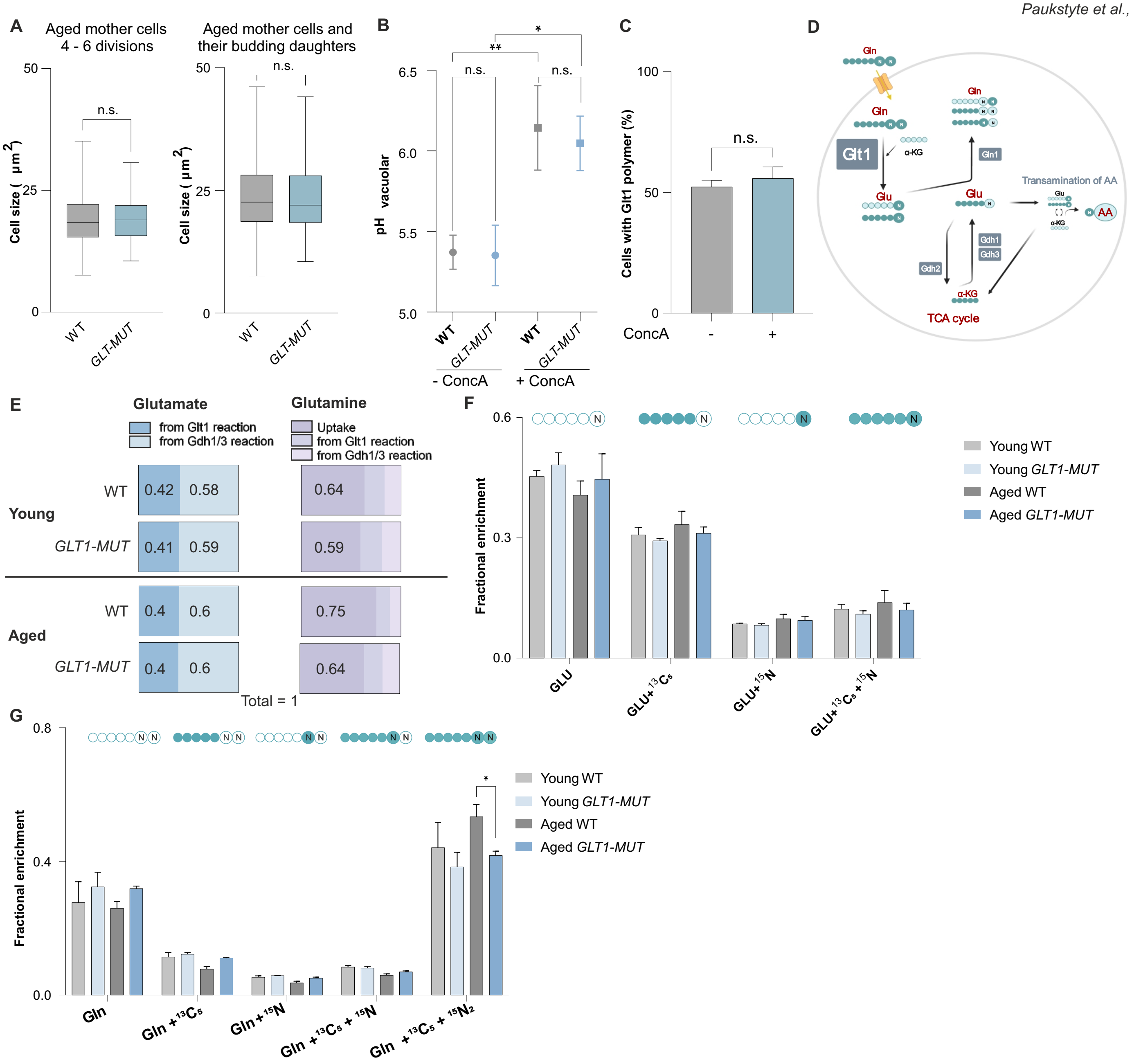

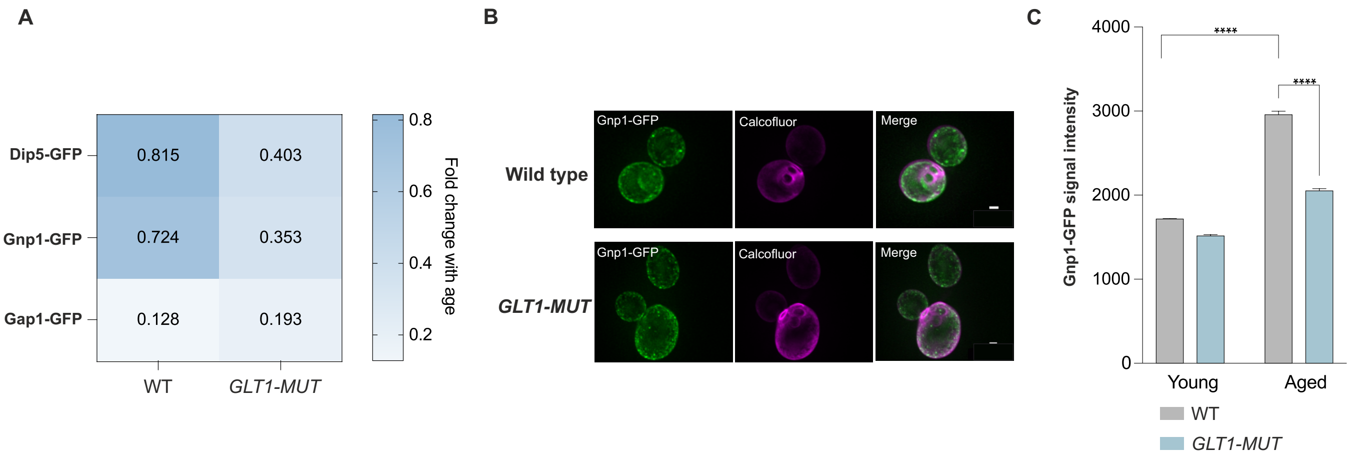

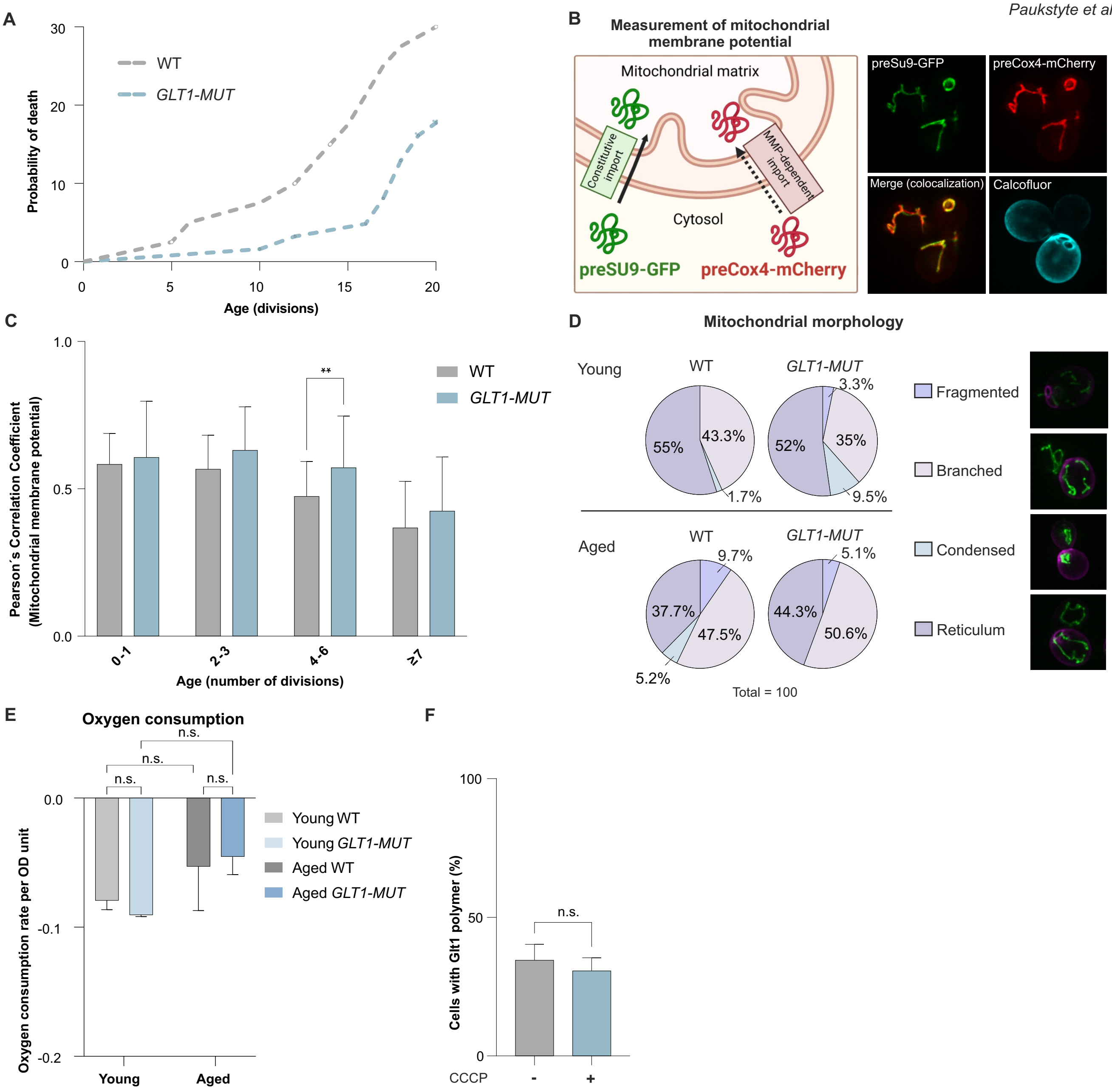
